# The yeast claudin Dcv1 is important for demarcating the front domain of pheromone-responding yeast cells

**DOI:** 10.1101/2022.04.18.488708

**Authors:** Madhushalini Sukumar, Reagan DeFlorio, Chih-Yu Pai, David E. Stone

## Abstract

Cell polarization in response to chemical gradients is important in development and homeostasis across eukaryota. Chemosensing cells orient toward or away from gradient sources by polarizing along a front-rear axis. Using the chemotropic mating response of the budding yeast *S. cerevisiae* as a model of environmentally-induced cell polarization, we found that Dcv1, a claudin homolog, is a determinant of front-rear polarity. Although Dcv1 localized uniformly on the plasma membrane (PM)^†^ of vegetative cells, it was confined to the rear of cells responding to pheromone, away from the pheromone receptor and mating projection. Deletion of *DCV1* conferred mislocalization of sensory, polarity, and trafficking proteins, as well as the PM lipids ergosterol, phosphatidylinositol-4,5-bisphosphate (PIP2), and phosphatidylserine (PS). These phenotypes correlated with defects in pheromone-gradient tracking and cell fusion. We propose that the novel claudin-like and rear-domain protein, Dcv1, demarcates the mating-specific front domain primarily by restricting PM lipid distribution. Consistent with this hypothesis, a mutation that blocks ergosterol biosynthesis partially phenocopies *dcv1*Δ.

**Summary statement:** The yeast claudin Dcv1 facilitates the proper localization of plasma membrane lipids and proteins that are required for front-rear polarity, efficient chemotropism, and cell fusion.

## INTRODUCTION

Cell polarization is likely essential for all species. In metazoans, cell polarity is integral to differentiation and development. The establishment of anterior-posterior polarity during embryogenesis, planar cell polarity in tissues, apico-basolateral polarity of epithelial cells, and front-to-rear polarity of migrating cells are examples of distinct polarity states established by cells for specialized functions.

Cell polarity can arise either spontaneously or in response to intrinsic or extrinsic cues. Directed cell migration (chemotaxis) and directed cell growth (chemotropism) are well-studied models of polarity induced by extrinsic cues. To move or grow in response to a directional signal, a chemosensing cell must solve three problems: First, it must interpret shallow and complex extracellular gradients to locate the chemoattractant source; second, it must align its axis of polarity toward the gradient source; third, it must generate distinct front and rear domains. In migrating cells, for example, signaling proteins are recruited to the front domain where they promote pseudopod formation at the leading edge, whereas opposing activities localize to the rear domain, or uropod, where they effect retraction of the lagging edge. The front and rear domains of migrating cells are also distinguished by their constituent lipids. For example, cholesterol accumulates in the plasma membrane at the leading edge and is required for polarization of phosphoinsositide 3-kinase, C-C chemokine receptor type 5, and proteins containing pleckstrin homology (PH) domains. Moreover, the PM of migrating cells exhibit a font-to-rear microviscosity gradient that depends on cholesterol polarization (Vasanji et al., 2004).

The unicellular eukaryote *Saccharomyces cerevisiae* has been extensively used to study cell polarity – both the intrinsically-regulated polarized growth of daughter cells (also called buds) during its vegetative cell cycle and the externally-regulated polarized growth of mating projections (also called shmoos) during the sexual reproduction stage of its lifecycle. The latter is a chemotropic process. Each of the two yeast haploid mating types, *MAT***a** and *MAT*α, secretes a peptide pheromone that activates a G-protein-coupled receptor (GPCR) on cells of the opposite type. The pheromone-bound receptor activates its cognate heterotrimeric G protein, causing Gα-GTP to dissociate from Gβγ. Free Gβγ then signals to the nucleus through a MAP kinase (MAPK) cascade, inducing transcription of mating-specific genes and cell cycle arrest in late G1. In mating mixtures, cells find and contact a partner by determining the direction of the most potent pheromone source and polarizing their growth (shmooing) toward it. When cells are treated with isotropic pheromone, however, they shmoo adjacent to their last division site – i.e., at the default polarity site (DS) where they would have budded next if not arrested in G1. Whether stimulated by isotropic pheromone or a pheromone gradient, the cell forms a front domain specialized for fusion with a mating partner.

As in migrating cells, the front domain of pheromone-stimulated yeast cells comprises specific PM lipids as well as proteins. Lipids that are enriched in the PM of the mating projection include ergosterol (Bagnat and Simons, 2002), phosphatidylserine (PS), and phosphatidylinositol-4,5-bisphosphate (PIP2) (Garrenton et al., 2010). Concentration of ergosterol in the mating projection is required for optimal GPCR signal transduction, downstream pheromone signaling, protein polarization, and mating (Bagnat and Simons, 2002; Garrenton et al., 2010; Jin et al., 2008; Morioka et al., 2013; Tiedje et al., 2007). PS anisotropy is required for the polarization and activation of the Rho GTPase Cdc42, an essential regulator of cell polarity, in both budding and pheromone-stimulated cells (Fairn et al., 2011; Sartorel et al., 2018). PIP2 membrane anisotropy is required for the shmoo-tip localization of the MAPK scaffold, Ste5, which presumably contributes to polarized MAPK activation (Garrenton et al., 2010). Another similarity between shmooing yeast and migrating cells is a front-to-rear differential in PM fluidity: the PM of the yeast mating projection is distinguished by high viscosity (Proszynski et al., 2006). Together, these observations suggest that PM lipid domains contribute to pheromone-induced front-rear polarity in mating yeast cells.

To search for novel regulators of front-rear polarity in pheromone-stimulated yeast cells, we conducted a directed screen for genes that affect polarization of the pheromone receptor. Here we characterize one such gene, the claudin homologue, *DCV1*. Claudins are a family of tetra-spanning, integral membrane proteins that are essential components of tight junctions and are required for the apico-basal polarity in epithelial cells. Consistent with its identification as a claudin-like protein (Martin et al., 2011), Dcv1 localized uniformly to the PM in vegetative cells. In pheromone-stimulated cells, however, Dcv1 localized anisotropically – away from the mating projection and receptor. Deletion of *DCV1* conferred defects in receptor polarization and orientation, shmoo morphology, and zygote formation; mislocalization of proteins involved in trafficking, cell polarity, and cell fusion; and mislocalization of PM lipids. We propose that Dcv1 facilitates the formation and/or maintenance of pheromone-induced front/rear membrane domains that are required for efficient chemotropism and mating.

## MATERIALS AND METHODS

### Molecular and microbiological techniques

Standard methods were used for microbial culture and molecular manipulation as described (Sherman et al., 1986; Ausubel et al., 1994; Guthrie and Fink, 2002). Yeast cultures were synchronized in G1 by centrifugal elutriation as described (Suchkov et al., 2010). Yeast and bacterial strains used in this study are listed in Supplementary Tables 3 and 4, respectively.

### Yeast strain construction

All yeast strains were derived from BY4741 (*MAT***a** *his3*Δ *leu2*Δ *met15*Δ *ura3*Δ), the strain used to construct the haploid yeast knockout collection (Brachmann et al., 1998). Hpa1-cut LHP1921 was integrated into strains MSY101, MSY116, and MSY288 to generate strains MSY128, MSY143, and MSY292 (*MAT**a** STE2-GFP*, *MAT**a** dcv1*Δ *STE2-GFP*, and *MAT**a** erg6*Δ *STE2-GFP),* respectively. MSB104 was transformed into strain MSY143 to generate strain MSY376 (*MAT**a** dcv1*Δ *STE2-GFP DCV1_[120]_-RFP*). MSB45 and MSB20 were transformed into strain MSY128 to generate strains MSY213 and MSY198 (*MAT**a** STE2-GFP* and *MAT**a** STE2-GFP GAL1*-*DCV1*), respectively. MSB59 was transformed into strains MSY101 and MSY116 to generate strains MSY351 and MSY352 (*MAT**a** GAL-GFP-PH^PLCδ^-PH^PLCδ^-GFP* and *MAT**a** dcv1*Δ *GAL-GFP-PH^PLCδ^-PH^PLCδ^-GFP*), respectively. MSB68 was integrated into strains MSY101 and MSY116 to generate strains MSY305 and MSY307 (*MAT**a** LACT-C2-GFP* and *MAT**a** dcv1*Δ *LACT-C2-GFP*), respectively. To switch their mating type to *MAT*α, strains MSY128 and MSY143 were transformed with MSB19 (p*GAL1-HO*). *HO* expression was induced with 2% galactose for 3 hours, after which cells were plated on synthetic glucose medium to isolate single colonies. *MAT*α cells were verified by their lack of response to alpha factor (αF, Genscript) and by their ability to mate with *MAT***a** cells. Cells confirmed to be *MAT*α were then grown on synthetic medium containing 5′ FOA to select for loss of MSB19, yielding strains MSY326 (*MAT*α) and MSY190 (*MAT*α *dcv1*Δ). Strain MSY190 was transformed with Bsu36I-cut DSB405 to RFP-tag *BUD1* in situ, generating strain MSY342 (*MAT*α *dcv1*Δ *RFP-BUD1*). BamH1-cut XWB087 was integrated into strains MSY128 and MSY143 to generate strains MSY378 and MSY380 (*MAT**a** STE2-GFP SLA1-RFP* and *MAT**a** dcv1*Δ *STE2-GFP SLA1-RFP*), respectively. The *URA3* coding sequence with flanking *SPA2* sequence (indicated by lowercase) was amplified from YCplac33 with the oligomers 5’ – atgggtacgtcaagcgaggtttctctcgcacatcatagaga tatcttccattactacgtcCCAGCTTTTCAATTCAATTC – 3’ and 5’ – ttacttcaacttcgaattcaaataatttattt cgtccttcaaacttgcctcttctacagtTTAGTTTTGCTGGCCGCATC – 3’. The resulting PCR product was used to transplace the native *SPA2* with *URA3* in strains MSY101 and MSY116, generating strains RDY321 and RDY333, respectively. *SPA2-GFP* was then amplified from RDB151 using the oligomers 5’ – atgggtacgtcaagcgaggt – 3’ and 5’ – ttagttttgctggccgcatcttctcaaatatgcttccc agcctgcttttctgtaaTTATTTGTATAGTTCATCCATGCCA – 3’, and the resulting PCR product was used to transplace *URA3* in strains RDY321 and RDY333 by selection on 5′ FOA, thereby generating strains RDY338 and RDY334 (*MAT**a** SPA2-GFP* and *MAT**a** dcv1Δ SPA2-GFP*), respectively. The *URA3* coding sequence with flanking *FUS1* sequence (indicated by lowercase) was amplified from YCplac33 with the oligomers 5’ – atggtagcaacaataatgcagacgacaacaactgtgctgacgacagtcgccgcaatgtctCCAGTTTTCAATTCAATTC – 3’ and 5’ – tcagtcgtattcttggagacagtcaccaggcacaatgcctctatcttcattgaggtatctTTAGTTTTGCTGGCCGCAT C – 3’. The resulting PCR product was used to transplace the native *FUS1* with *URA3* in strains MSY101 and MSY116, generating strains RDY363 and RDY364, respectively. *FUS1-GFP* was then amplified from 379 using the oligomers 5’ – atggtagcaacaataatgcag – 3’ and 5’ – ttagttttgctggccgcatcttctcaaatatgcttcccagcct gcttttctgtaaTTATTTGTATAGTTCATCCATGCCA – 3’, and the resulting PCR product was used to transplace *URA3* in strains RDY363 and RDY364 by selection on 5′ FOA, thereby generating strains RDY365 and RDY367 (*MAT**a** FUS1-GFP* and *MAT**a** dcv1Δ FUS1-GFP*), respectively. All genomic modifications were verified by DNA sequencing (University of Illinois at Chicago, Research Resource Center, Sequencing core).

### Plasmid construction

The first 360 bases of the *DCV1* coding region and 400 bases upstream were amplified using the oligomers 5’ – ATGCGGATCCTTTGTACAATTCATCCATACCATGGG – 3’ and 5’ – GCATAAGCTTTCTTGAGATGGGCGTTCG – 3’, and the resulting PCR product digested with HindIII and SalI. The oligomers 5’ – GCATAAGCTTTCTTGAGATCGGGCGTTCG – 3’ and 5’ – GATCGAGCTCGGTCTGTGGCAATGTTTGTC – 3’ were used to amplify the remainder of the *DCV1* coding sequence and 300 bases of downstream flanking sequence, and the resulting PCR product digested with BamHI and SstI. RFP was amplified from DSB405 using the oligomers 5’ – ATCGATCGGTCGACATGGTGAGCAAGGGCGAGGAGG – 3’ and 5’– ATCGGGATCCCTTGTACAGCTCGTCCATGCCGTACAGG – 3’, and the resulting PCR product digested with SalI and BamHI. The digested PCR products were sequentially ligated into YCplac33 to generate MSB104 (DCV1_[120] -_RFP). The EcoRI- and BamHI-cut GAL1/GAL10 promoter from ZWE159 was subcloned into YIplac211 to generate MSB67 (Yiplac211-GAL1/GAL10). The Lact-C2 domain was amplified from MSB56 using the oligomers 5’ – GCAGACGGATCCGCCACCATGGTGAGCAAGGGCGAGG – 3’ and 5’ – GCAGACAAGCTTCTAACAGCCCAGCAGCTCCACTCG – 3’, and the PCR product ligated into MSB67 after digestion with BamHI and HindIII to generate MSB68 (GAL1-inducible PS reporter). All subclones were verified by DNA sequencing (University of Illinois at Chicago, Research Resource Center, Sequencing core).

### Cell staining

F-actin staining was performed as previously described (Pringle et al., 1989) using Alexa Fluor 594 Phalloidin (Invitrogen). For the trypan blue exclusion assay, log-phase cells were stained with 0.2% trypan blue for 15 minutes at room temperature, then washed twice with 1X PBS, spotted on slides, and scored under a phase contrast microscope at 40X magnification. Representative fluorescent images were deconvolved using Huygens Essential software (Scientific Volume Imaging) in standard mode. The filipin staining procedure was modified from the protocol described in Beh and Rine (2004). Briefly, log-phase cells were treated with 600 nM αF and 200µl aliquots were taken every 15 minutes. Cells were pelleted at 4000Xg and resuspended in 100µl of 1X PBS containing 4 µl of filipin (Sigma) stock solution (5 mg/ml in ethanol). Cells were spotted onto slides and images were acquired for 2 sec using a Zeiss observer Z.1 microscope, DAPI filter, 100X oil immersion objective, and Zeiss Zen software. 16 Z-sections 0.25 µm apart were acquired around the center slice of each cell at each time point and deconvolved using Zeiss Zen software.

### Drug treatment

To assay sensitivity to PM and cell wall stressors, WT and *dcv1*Δ cells were grown to log-phase in rich medium; the Dcv1-overexpression and corresponding control cells were grown to log-phase in synthetic medium containing either 2% dextrose (uninduced Dcv1) or 2% galactose (induced Dcv1). A series of aliquots, starting with 3 X 10^6^ cells and decreasing by factors of 10, were spotted onto the appropriate plate medium (dextrose or galactose) containing either Congo red (Sigma; 100 µg/ml), SDS (Sigma; 0.001%), Caffeine (Sigma; 12 mM), Hygromyocin B (50 µg/ml), Ethanol (4%), Sodium chloride (Sigma; 0.4 M), or no stressor. The plates were incubated at 30°C for two overnights before photographing.

### Fluorescent imaging of pheromone-treated cells

Log-phase cultures were synchronized in G1 and treated with 600nM αF. 200 µl aliquots were taken every 15 minutes, pelleted at 4000Xg, and resuspended in 1X PBS. To visualize Ste2-GFP, cells were spotted onto slides and images were acquired for 8 sec using an Axioscop 2 microscope (Carl Zeiss), FITC filter, 100X oil immersion objective, and Axiovision software. To co-visualize Ste2-GFP and Dcv1_[120]_-RFP, cells were spotted onto slides and images were acquired using a Deltavision Elite microscope (GE Healthcare Biosciences) with a 60X oil immersion objective, a Front Illuminated sCMOS camera, and SoftWoRx software. Ste2-GFP was imaged at 461-489 nm for 200 ms using 10% maximum intensity. Dcv1_[120]_-RFP was imaged at 529-556 nm for 1 sec using 50% maximum intensity. 5 z-sections 0.5 µm apart were acquired around the center slice of each cell at each time point. For lipid reporter localization studies, cells were grown to log-phase in synthetic 2% sucrose medium. Reporter expression was induced with 2% galactose for 1 hr. Cells were then spun down, resuspended in synthetic 2% dextrose medium to turn off GAL induction, and spread at a density of 14,000 cells/mm^2^ on 1% agarose pads made from synthetic medium containing 1.8 µM αF. The pads were maintained at 30°C for the duration of the experiment. Images were acquired at 10-minute intervals using the Deltavision Elite microscope and 60X objective described above. 16 z-sections 0.25 µm apart were acquired around the center slice of each cell at each time point. The PIP2 reporter was imaged at 461-489 nm for 100 ms using 10% maximum intensity. The PS reporter was imaged at 461-489 nm for 400 ms using 10% maximum intensity. Representative fluorescent images were deconvolved using Huygens Essential software (Scientific Volume Imaging) in standard mode. DIC center-slice reference images were acquired at 100% of maximum intensity using polarized light for 5 ms.

### Time lapse imaging of cells in mating mixtures

Time lapse analysis of mating mixtures was performed as described (Wang et al., 2019). The indicated *MAT***a** and *MAT*α strains were grown to mid-log phase in synthetic 2% dextrose medium at 30°C, mixed 1:1, and spread at a density of 14,000 cells/mm^2^ on agarose pads made from synthetic dextrose medium. Mating mixtures were maintained at 30°C using a DeltaVision environment control chamber except as noted below. Images were acquired from 15 fields at 5- or 10-minute intervals using a DeltaVision Elite Deconvolution Microscope (GE Healthcare Biosciences) with a 60x oil immersion objective and a Front Illuminated sCMOS camera. 16 z-sections 0.25 µm apart were acquired around the center slice of each cell at each time point. GFP was imaged at 461-489 nm for 150 ms (Spa2-GFP and Fus1-GFP) or 200 ms (Ste2-GFP) using 10% maximum intensity. RFP was imaged at 529-556 nm for 150 ms (Bud1-RFP) or 200 ms (Sla1-RFP) using 10% maximum intensity. Representative fluorescent images were deconvolved using Huygens Essential software in standard mode. DIC images were acquired at 100% of maximum intensity using polarized light for 5 ms.

### Image analysis

Cells were randomly chosen for analysis. In mating mixtures, only *MAT***a** cells that fused with a partner were analyzed. Cell fusion was detected by the movement of Bud1-RFP from the *MAT*α partner to the *MAT***a** partner (Figs. 4, 6-8), or by the movement of Sla1-RFP from the *MAT***a** partner to the *MAT*α partner (Fig. 5). For Spa2-GFP analysis (Fig. 6), only cells that exhibited clear bud neck localization and had finished the cell cycle were scored. Crescent size, cell size, zygote bud position, and pixel intensities were determined using the ImageJ (National Institutes of Health) segmented line tool set at a width of 2 pixels. The ImageJ Look-Up-Tables fire tool was used to identify the demarcation between inner and outer crescents and to determine the ends of crescents. Crescent size was calculated as a percentage of the total cell circumference. The size of tracking receptor crescents was measured at the midpoint between the DS and the chemotropic growth and fusion site (CS). All angles were measured using the ImageJ angle tool. Prezygotes were identified based on two criteria: no visible cell walls between the two partners in the region of contact; Ste2-GFP tightly localized as a bar at the fusion zone. Mating efficiency indicates the number of observed zygotes as a percentage of the potential zygotes. The number of potential zygotes was determined by counting the number of *MAT**a*** cells initially positioned less than 3 µm away from a *MAT*α cell.

### Statistical quantifications

GraphPad Prism 8 was used for all graphical representations and statistical calculations. The *p*-values for all comparisons excluding percentages were determined by two-tailed unpaired t-tests. The *p*-values for comparisons of percentages were determined by chi-square test.

## RESULTS

### Dcv1 affects pheromone-induced receptor and actin-cable polarization but not receptor endocytosis

To identify novel regulators of pheromone-induced front-rear polarity, we conducted a directed genetic screen using polarization of the pheromone receptor to the mating projection as a proxy. Because mating is a haploid-specific process, and because the pheromone receptors are only expressed in haploid cells, we compiled a list of haploid-specific genes (Table S1) from which we selected candidate regulators based on their published involvement in signaling or cell polarity and/or their pheromone-dependent expression. The GPCR expressed by *MAT***a** cells, *STE2,* was tagged with GFP in situ in the corresponding deletion strains (Brachmann et al., 1998). The intracellular localization of the receptor was then imaged in each of the resulting strains before and after pheromone treatment of G1-synchronized cultures. As previously shown (Suchkov et al., 2010), pheromone induced global internalization of Ste2-GFP from the PM of wild type (WT) cells, after which Ste2-GFP reappeared as polarized PM crescents just before morphogenesis (Fig. 1A). Deletion of *DCV1* conferred a significant increase in cell size (Fig. S1) and a defect in receptor polarization (Fig. 1A). Although the kinetics of pheromone-induced receptor internalization and formation of polarized receptor crescents were indistinguishable in WT and *dcv1*Δ cells (Fig. 1B,C), mean receptor crescent size was dramatically increased in the absence of Dcv1 (Fig. 1D). Whereas the receptor crescents spanned about one third the circumference of WT cells and had clearly defined boundaries, the receptor crescents in *dcv1*Δ cells comprised a higher-signal region spanning about one third of the cell (hereafter called the inner crescents) flanked by lower-signal regions that gradually disappeared about halfway around the cell (hereafter called the total crescents) (Fig. 1A,D). Unlike the receptor crescents in WT cells, the crescents in *dcv1*Δ cells lacked a clear boundary.

**Figure 1.**
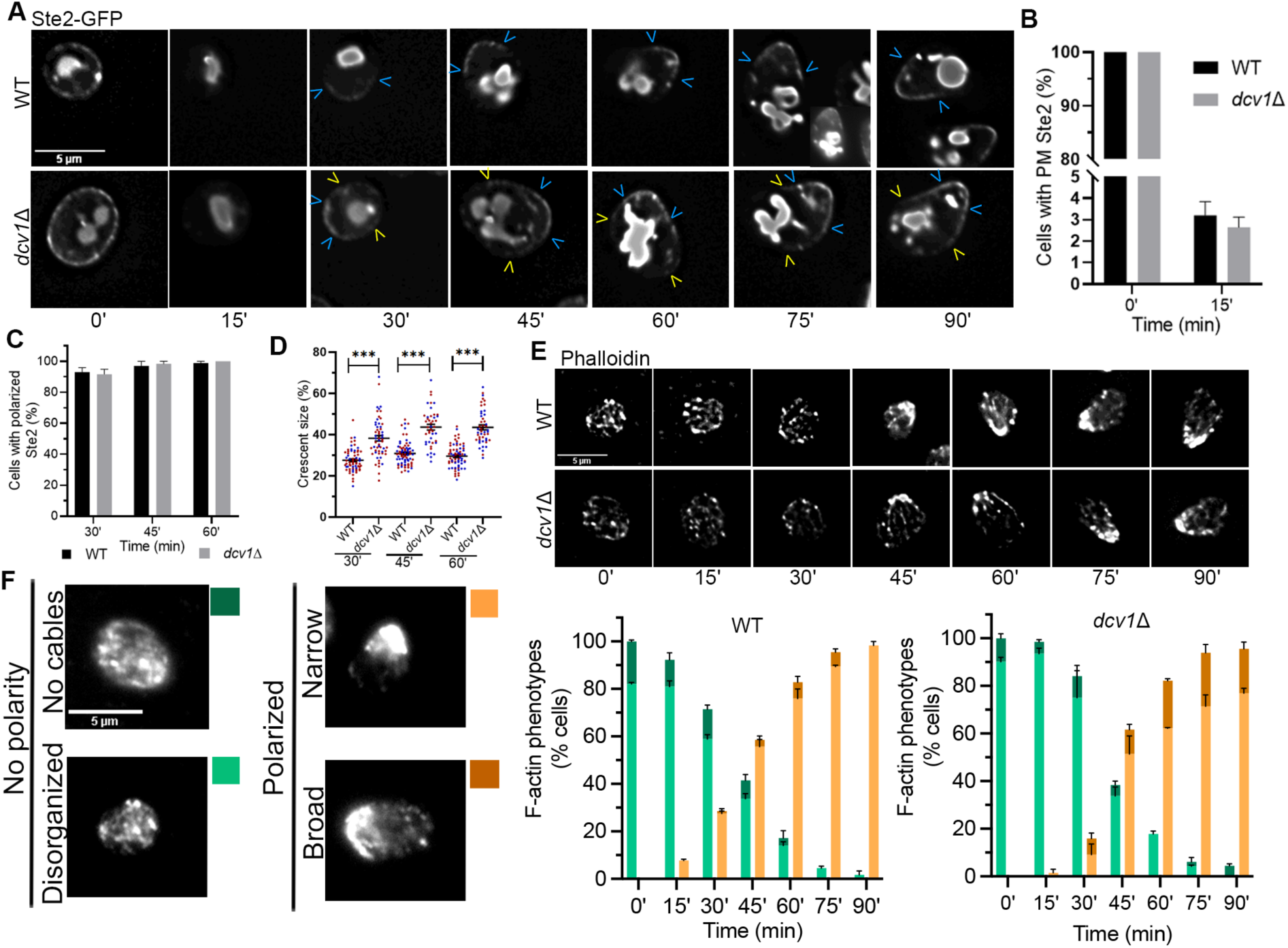
*dcv1*Δ confers a defect in pheromone-induced receptor polarization. (A-D) G1-synchronized daughter cells were isolated by centrifugal elutriation, treated with 600 nM pheromone, and imaged every 15 minutes. (A) The receptor is less well polarized in *dcv1*Δ cells. Representative images of WT and *dcv1*Δ cells expressing Ste2-GFP are shown. Blue arrowheads indicate the edges of inner crescents; yellow arrowheads indicate the edges of total crescents. (B) Receptor internalization kinetics are similar in WT and *dcv1*Δ cells. Bar graphs represent the mean percentage of cells with detectable Ste2-GFP on the PM ± s.e.m. at the indicated time points measured in two independent experiments (WT: 0’ = 100 ± 0, 15’ = 3.2 ± 0.6; *dcv1*Δ: 0’ = 100 ± 0, 15’ = 2.7 ± 0.5. n: WT = 43; *dcv1*Δ = 85.). (C) Receptor polarization kinetics are similar in WT and *dcv1*Δ cells. Bar graphs represent the mean percentage cells with polarized Ste2 ± s.e.m. at the indicated time points, measured in two independent experiments (WT: 30’ = 93.0 ± 2.8, 45’ = 97.0 ± 3.0, 60’ = 98.8 ± 1.2; *dcv1*Δ: 30’ = 91. 7 ± 3.2, 45’ = 98.3 ± 1.7, 60’ = 100 ± 0. n: WT = 43; *dcv1*Δ = 85.). (D) *dcv1*Δ cells form significantly larger receptor crescents. The size of total receptor crescents as a percentage of cell circumference was determined at the indicated times after pheromone treatment. Data points represent crescent sizes measured in two independent experiments, indicated by color. Horizontal lines and error bars indicate the means ± s.e.m. (WT: 30’ = 27.6 ± 0.7, 45’= 30.8 ± 0.7, 60’= 29.6 ± 0.7; *dcv1*Δ: 30’ = 38.2 ± 1.3, 45’ = 43.6 ± 1.4, 60’ = 43.5 ± 1.2. n: WT = 70; *dcv1*Δ = 48. ***p < 0.0001). (E) F-actin polarizes to the tip of the mating projection in WT and *dcv1*Δ cells. Images are representative of cells treated with pheromone and stained with 1.5 µM phalloidin at the indicated time points. (F) F-actin polarization dynamics are similar in WT and *dcv1*Δ cells. Cells were classified according to their F-actin phenotypes. Representative images illustrating the categories are shown: no cables (dark green); disorganized F-actin (light green); narrowly polarized F-actin (orange); broadly polarized F-actin (brown). Bar graphs represent the mean percentage of cells in each category ± s.e.m. at the indicated time points measured in two independent experiments (mean ± s.e.m. values in Table S2A. n *≥* 46).

Although pheromone-induced receptor polarity does not depend on actin-cable directed secretion, F-actin cables do contribute to its maintenance and amplification (Ayscough and Drubin, 1998; Ismael et al., 2016; Suchkov et al., 2010). Therefore, the receptor depolarization we observed in *dcv1*Δ cells could be due to a defect in directed secretion. To test this possibility, we visualized actin cables in G1-synchronized WT and *dcv1*Δ cells by staining with Phalloidin before and after pheromone treatment. Cells were scored as displaying one of the following phenotypes: no cables, disorganized cables, narrowly polarized cables, and broadly polarized cables. We found that the kinetics of pheromone-induced cable polarization were similar in the control and experimental strains; however, the incidence of broadly polarized cables, which correlated with broader mating projections, was higher in the *dcv*1Δ cells (Fig. 1E,F). Because the receptor affects the actin cytoskeleton via the binding of its G*βγ* protein to Far1-Cdc24-Cdc42 (Butty et al., 1998; Nern and Arkowitz, 1998; Nern and Arkowitz, 1999), depolarization of the receptor in *dcv1*Δ cells could result in a corresponding broadening of cable polarity. Alternatively, the absence of Dcv1 could adversely affect cable polarity, resulting in depolarization of the receptor. Notably, receptor polarization was detectable prior to actin polarization in most *dcv1*Δ cells (Fig. 1C,F), suggesting that Dcv1 affects receptor polarity independently of actin-cable directed secretion.

### Dcv1 localizes uniformly to the plasma membrane of vegetative cells and away from the receptor crescent in shmooing cells

Dcv1 is predicted to be a 4-pass integral membrane protein and a member of the claudin superfamily (Martin et al., 2011). To visualize Dcv1 in live cells and compare its localization to that of the receptor, we constructed an RFP-tagged *DCV1* (Dcv1_[120]_-RFP) reporter and expressed it in a *dcv1*Δ *STE2-GFP* strain. Dcv1_[120]_-RFP colocalized with Ste2-GFP on the PM of vegetative cells, and like the receptor reporter, it transiently disappeared from the PM in pheromone-treated cells. After recovery of the PM reporter signals and just before morphogenesis, the receptor and Dcv1 localized inversely to one another in a majority of the cells: As the receptor polarized to the mating projection, Dcv1 concentrated at the rear (Fig. 2).

**Figure 2.**
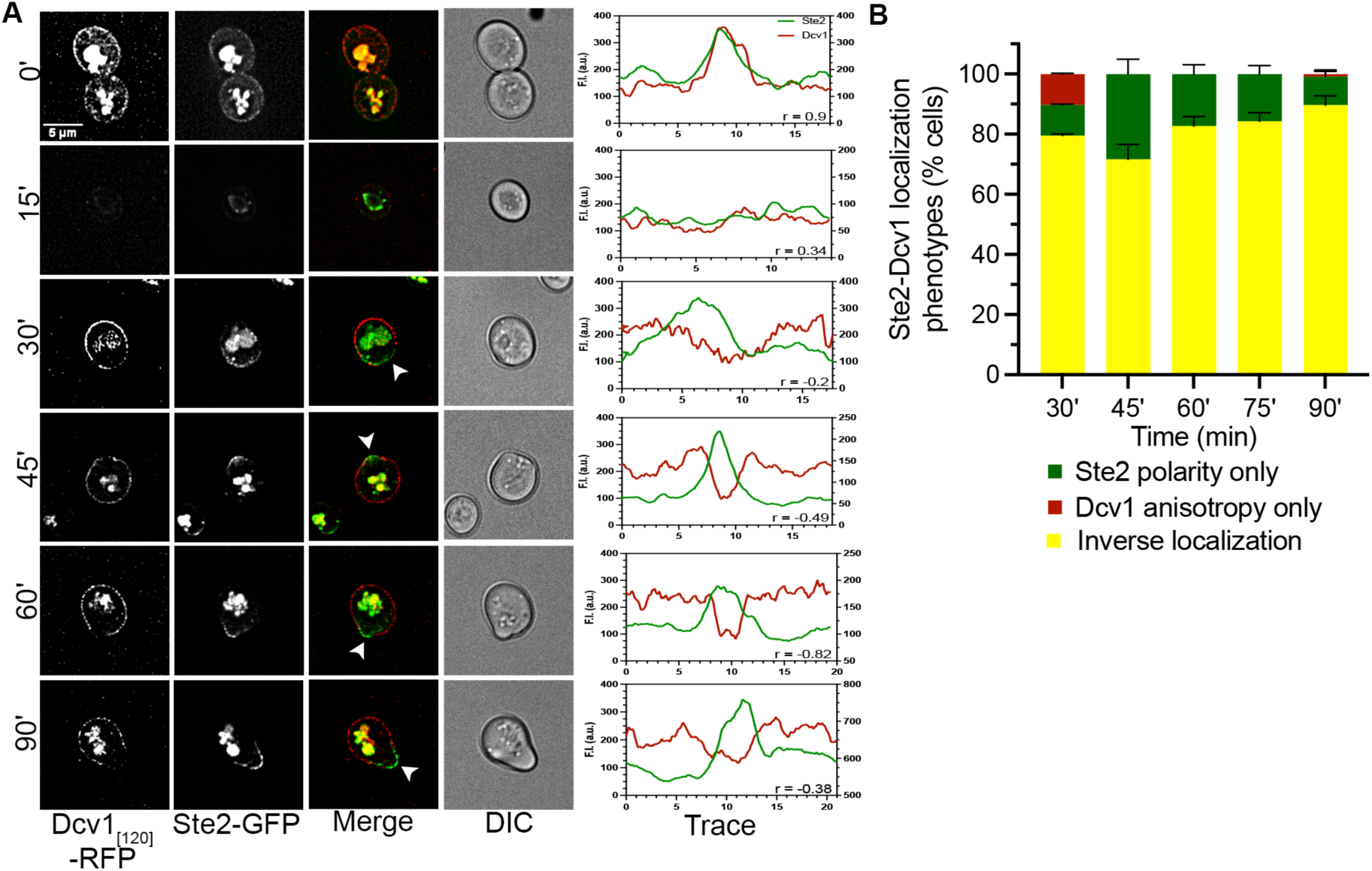
Dcv1 and the receptor inversely localize in pheromone-treated cells. Log-phase cells were treated with 600 nM pheromone and imaged every 15 minutes. (A) Representative images of cells co-expressing Dcv1_[120]_-RFP and Ste2-GFP at the indicated time points are shown. PM regions where the receptor polarizes and the Dcv1 signal is undetectable are indicated by white arrowheads. Line graphs show the distribution of each reporter on the PM (four-point rolling average). Green line, Ste2-GFP; red line, Dcv1_[120]_-RFP; r values, Pearson’s correlation coefficient. The left and right y-axes correspond to the Ste2-GFP and Dcv1_[120]_-RFP fluorescent intensity (F.I.) values in arbitrary units, respectively. The bottom cell was traced in the 0’ image. (B) Inverse localization of receptor and Dcv1 in cell populations. Cells with detectable Dcv1 PM signal were classified according to Ste2 and Dcv1 localization as follows: polarized Ste2, uniform Dcv1 (green); anisotropic Dcv1, depolarized Ste2 (red); polarized Ste2, anisotropic Dcv1 (yellow). Bar graphs represent the mean percentages of cells in each category ± s.e.m. measured at the indicated time points in two independent experiments (mean ± s.e.m. values in Table S2B. n *≥* 70).

### Dcv1 affects cell integrity and the distribution of plasma membrane lipids

We next examined the effects of Dcv1 overexpression on receptor polarity. Although we could detect no effect on the receptor, cells overexpressing Dcv1 were abnormally prone to lyse. To confirm this observation, we used the trypan blue (TB) assay for cell viability. TB stains dead cells blue, whereas viable cells with an intact PM exclude the dye. Dcv1 overexpression increased the percentage of TB-stained cells five-fold, consistent with a defect in cell integrity (11.1 ± 0.7% Dcv1-overexpressing cells vs. 2.1 ± 0.2% control cells; n = 900 in three independent experiments; *p* = 0.0001). To further explore the connection between Dcv1 and cell integrity, we spotted Dcv1-overexpressing cells and *dcv1*Δ cells on medium containing various cell wall and PM stressors. Overexpression of Dcv1 increased sensitivity to Congo red, hygromyocin B, SDS, ethanol and NaCl (Fig. S2A); *dcv1*Δ conferred a clear hypersensitivity to congo red and a slight increase in sensitivity to caffeine (Fig. S2B). These data demonstrate that both excess and absence of Dcv1 adversely affect cell integrity.

In higher eukaryotes, the misregulation of claudins confers a loss of cell polarity and integrity, and these effects are correlated with altered membrane domains (Lingaraju et al., 2015). To test the possibility that Dcv1 also influences membrane domains, we assayed the effect of *dcv1*Δ on the polarization of sterols, PIP2, and PS in both vegetative and pheromone-treated cells.

To assay sterol localization, we stained cells with filipin dye. The filipin staining patterns of vegetative WT and *dcv1*Δ cells were indistinguishable: about 60% of each population showed a uniform distribution of sterols on the PM while the remainder exhibited a slight asymmetry. In contrast, *dcv1*Δ cells showed a defect in pheromone-induced sterol polarization (Fig. 3A-C). Whereas most WT cells polarized sterols to a single, well-focused spot on the PM before the initiation of shmooing (morphogenesis), there was a high fraction of cells with two or more polarized sterol spots in the *dcv1*Δ culture (Fig. 3B). The incidence of *dcv1*Δ cells with multiple spots decreased after morphogenesis, perhaps due to the directed delivery of vesicles to the growth site; however, the percentages of mutant cells with aberrantly polarized sterols were significantly higher than in the control culture at all time points (Fig. 3B,C). These results indicate that *dcv1*Δ confers a defect in coalescing sterols to a single site on the PM prior to the onset of polarized growth.

**Figure 3.**
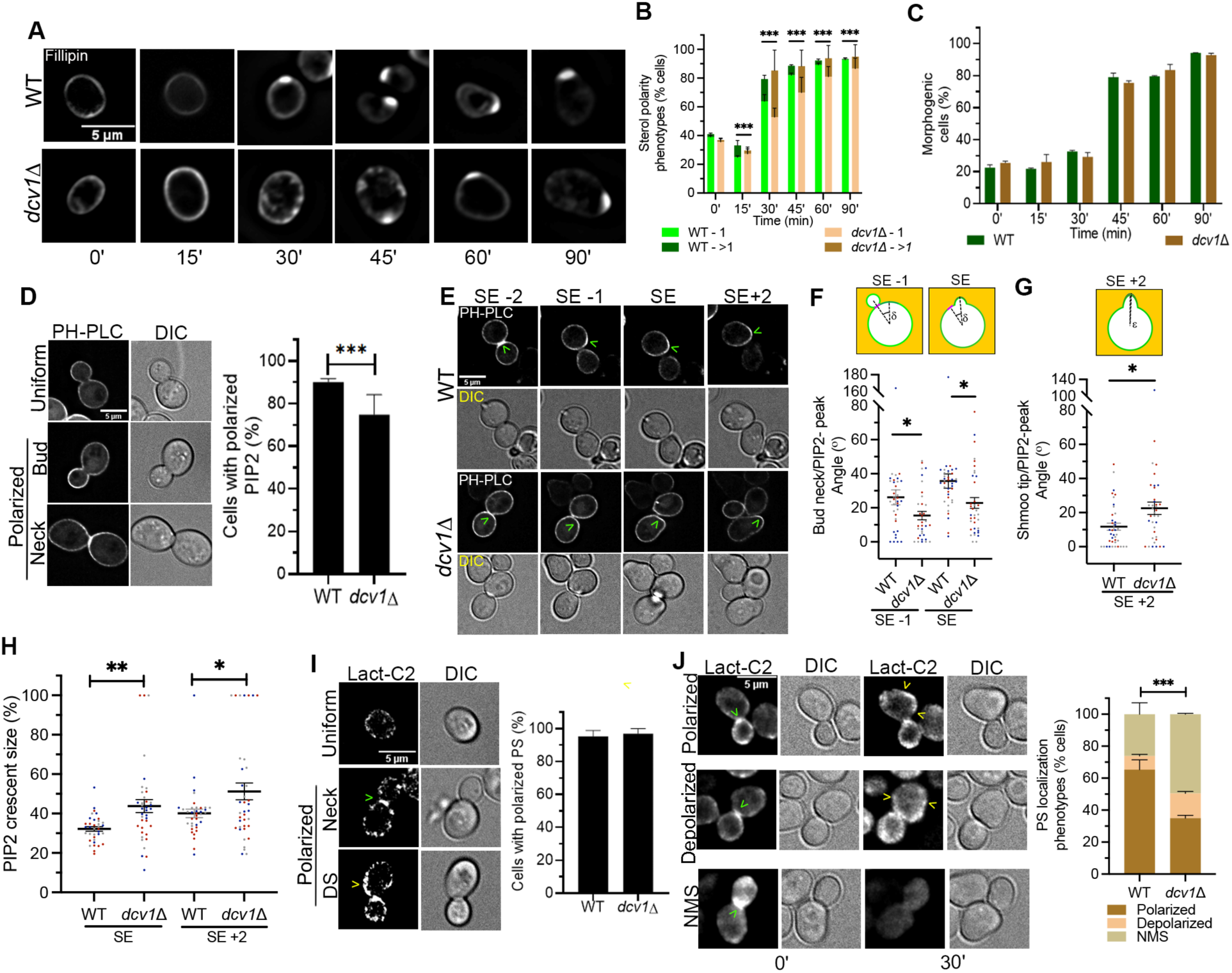
Cells lacking Dcv1 exhibit altered PM lipid distribution. (A-C) Log-phase WT and *dcv1*Δ cells were G1-synchronized and treated with 600 nM pheromone. Samples were stained with 305 µM filipin to visualize sterols at the indicated time points. (A) *dcv1*Δ confers aberrant sterol polarization. Center slices of representative images are shown. (B) Quantification of sterol patches in polarized cells. Polarized Cells were classified as having either one polarized sterol patch or more than one polarized sterol patch at the indicated time points. Bar graphs represent the mean percentage of cells in each category ± s.e.m. measured in two independent experiments (mean ± s.e.m. values in table S2C. n *≥* 40. \*\*\**p*_[chi sq]_ ≤ 0.0001). (C) The kinetics of morphogenesis are similar in WT and *dcv1*Δ cells. Cells were characterized based on their shape and the percentage of polarized cells were determined at the indicated time points. Bar graphs represent the mean percentage of cells ± s.e.m. measured in two independent experiments (mean ± s.e.m. values in Table. S2D. n *≥* 40). (D) *dcv1*Δ decreases the proportion of vegetative cells that exhibit polarized PIP2 localization. Representative images of log-phase WT and *dcv1*Δ cells expressing the GAL-induced PIP2 reporter are shown. Cells were scored as exhibiting either uniform or polarized PIP2 localization. Bar graphs represent the mean percentage of polarized cells ± s.e.m. measured in three independent experiments (WT = 90.0 ± 1.7; *dcv1*Δ = 74.8 ± 9.3. n *≥* 44. \*\*\**p*_[chi sq]_ = 0.0001.). (E-H) PIP2 localization in pheromone-treated cells. WT and *dcv1*Δ cells expressing the PIP2 reporter were exposed to 3 µM pheromone on agarose pads and imaged every 15 minutes. SE-2, two time points before shmoo emergence; SE-1, one time point before shmoo emergence; SE, shmoo emergence; SE+2, two timepoints after shmoo emergence. (E) Pheromone-treated *dcv1*Δ cells exhibit PIP2 polarization defects. Center slices of representative images are shown. Green open arrowheads point to peak of PIP2 polarity. (F) *dcv1*Δ cells are defective in switching PIP2 polarity from the neck to the presumptive DS. The angle from the center of the neck to the peak of the PIP2 signal was measured one time point before shmoo emergence and at shmoo emergence as illustrated in the diagrams. δ is the angle formed by rays drawn from the middle of the cell to the center of the neck and to the peak of PIP2 polarity. Data points represent the neck/PIP2-peak angle measured in three independent experiments, indicated by color. Horizontal lines and error bars represent the mean ± s.e.m. (WT: SE-1 = 26.1 ± 4.4, SE = 35.7 ± 4.2; *dcv1*Δ: SE-1 = 15.4 ± 2.5, SE = 22.8 ± 3.2. n = 38. \**p* ≤ 0.04). (G) *dcv1*Δ cells are defective in aligning PIP2 polarity with the shmoo tip. The angle from the shmoo tip to the peak of PIP2 polarity was measured in mature shmoos (SE+2) as illustrated in the diagram. ε is the angle formed by rays drawn from the middle of the cell to the shmoo tip and to the peak of PIP2 polarity. Data points represent the shmoo tip/PIP2-peak angle measured in three independent experiments, indicated by color. Horizontal lines and error bars represent the mean ± s.e.m. (WT = 11.8 ± 2.1; *dcv1*Δ = 22.5 ± 3.7. n = 38. \**p* = 0.01). (H) PIP2 crescent sizes at SE and SE+2. Data points represent crescent sizes as a proportion of cell circumference measured in three independent experiments, indicated by color. Lines and error bars represent the mean ± SEM. (WT: SE = 32.2 ± 1.1, SE+2 = 40.1 ± 2.1; *dcv1*Δ: SE = 43.8 ± 3.3, SE+2 = 51.2 ± 4.3. n = 38. \**p* = 0.025; \*\**p* = 0.0017). (I) PS localizes to sites of polarized growth in vegetative cells. Representative images of log-phase WT and *dcv1*Δ cells expressing the GAL- induced PS reporter and exhibiting either uniform or polarized PS are shown. The green open arrowhead points to bud neck localization and the yellow open arrowhead points to PS polarization at the DS. Bar graphs represent the mean percentage of polarized cells ± s.e.m. measured in two independent experiments (WT = 95.3 ± 3.6; *dcv1*Δ = 96.9 ± 3.1. n *≥* 48). (J) Pheromone-treated *dcv1*Δ cells exhibit defects in PS polarization. WT and *dcv1*Δ cells expressing the PS reporter were exposed to 3 µM pheromone on agarose pads. Quantification of PS polarization in pheromone-treated WT and *dcv1*Δ cells. The PS localization phenotype was classified as polarized (≤ 40% of the cell circumference, brown); depolarized (> 40% of the cell circumference, salmon); or showing no membrane signal (NMS, light brown). Center slices of representative images are shown. The green open arrowheads point to PS neck polarity and the yellow open arrowheads mark the edges of PS crescents. Bar graphs represent the mean percentage of cells in each category ± s.e.m. from two independent experiments (WT: polarized = 65.3 ± 6.1, depolarized = 8.5 ± 1.0, NMS = 26.2 ± 7.1; *dcv1*Δ: polarized = 35.0 ± 1.7, depolarized = 15.0 ± 1.2, NMS = 49.5 ± 0.5. n *≥* 65. \*\*\**p*_[chi sq]_ ≤ 0.0001).

To assay PIP2 localization, we imaged cells expressing the GFP-2xPH-PLCδ-GFP reporter (Stefan et al., 2002). In 90% of the budding WT cells, the PIP2 reporter concentrated in regions of polarized secretion – the PM of daughter cells and the mother-daughter neck (hereafter, the neck) where cytokinesis occurs – consistent with previous findings (Garrenton et al., 2010); the remaining 10% showed uniform PM localization. The PIP2 reporter also localized to regions of polarized secretion in vegetative *dcv1*Δ cells, although uniform distribution was seen in a significantly higher proportion of the mutant culture (Fig. 3D). In most WT cells responding to pheromone, the peak PIP2 signal moved from the neck to the presumptive DS prior to morphogenesis, forming a polarized crescent that remained centered around the elongating mating projection (Fig. 3E-G). In contrast, the peak PIP2 signal did not move from the neck to the DS before morphogenesis in most of the *dcv1*Δ cells, and this phenotype correlated with misalignment of the PIP2 crescents and shmoo tips (Fig. 3E-G). We also found that the mean size of the polarized PIP2 crescents was significantly larger in *dcv1*Δ shmoos than in WT shmoos (Fig. 3H).

To assay PS localization, we imaged cells expressing a galactose-inducible Lact-C2-GFP reporter (Yeung et al., 2008). In both vegetative WT and *dcv1*Δ cells, the PS reporter concentrated in regions of polarized secretion – the neck and the presumptive DS – consistent with previous findings (Fairn et al., 2011); only ∼5% of the cells showed uniform PM localization (Fig. 3I). In the majority of pheromone-treated WT cells, the PS reporter was enriched in the PM of the mating projection, whereas about a quarter of the WT cells exhibited no PM signal. In contrast, a significantly smaller fraction of *dcv1*Δ cells exhibited PS polarization and a significantly higher fraction showed no PM signal (Fig. 3J). The lack of membrane signal was not attributable to a reduction in reporter signal strength as the mean fluorescence intensity of the cytoplasm was comparable between the WT and mutant cells (Fig. S3).

### Dcv1 contributes to receptor polarization and orientation, and cell fusion during mating

In *MAT***a** *dcv1*Δ cells treated with isotropic pheromone, we observed depolarization of the *α*-factor receptor, Ste2, and defects in the localization of PM lipids. Because *MAT***a** cells that cannot polarize the *α*-factor receptor exhibit defects in gradient tracking and orientation toward mating partners (Ismael et al., 2016; Wang et al., 2019), and because the lipids whose proper localization depends on Dcv1 have established roles in pheromone signaling and zygote formation, we wondered whether Dcv1 contributes to polarized mating functions.

As a first test of this possibility, we compared time-lapse images of unilateral mutant (*MAT***a** *dcv1*Δ Ste2-GFP X *MATα DCV1*) and WT control (*MAT***a** *DCV1* Ste2-GFP X *MATα DCV1*) mating mixtures. The degree of receptor polarity was measured in *MAT***a** cells as their receptor crescents tracked upgradient toward their eventual mating partners and at the prezygote (PZ) stage, when they had established stable contact with their partners but had not yet begun to fuse. Only cells that ultimately formed zygotes (Fig. S4) were scored in this analysis. During both the gradient tracking and PZ stages, WT *MAT***a** cells mating with WT *MATα* cells displayed sharply demarcated receptor crescents that averaged about one third of their circumferences (Fig. 4A-C); these crescents invariably centered around the eventual fusion sites (Fig. 4D). Like *MAT***a** *dcv1*Δ cells treated with isotropic pheromone, those mating with WT *MATα* cells displayed significantly larger receptor crescents than the control cells — means of 1.8-fold and 1.56-fold larger during the tracking and PZ stages, respectively (Fig. 4A-C); these crescents often failed to center around the eventual fusion sites (Fig. 4D). The mutant cells also displayed significantly higher mean Ste2-GFP signals within the crescent and around PM (Fig. 4E).

**Figure 4:**
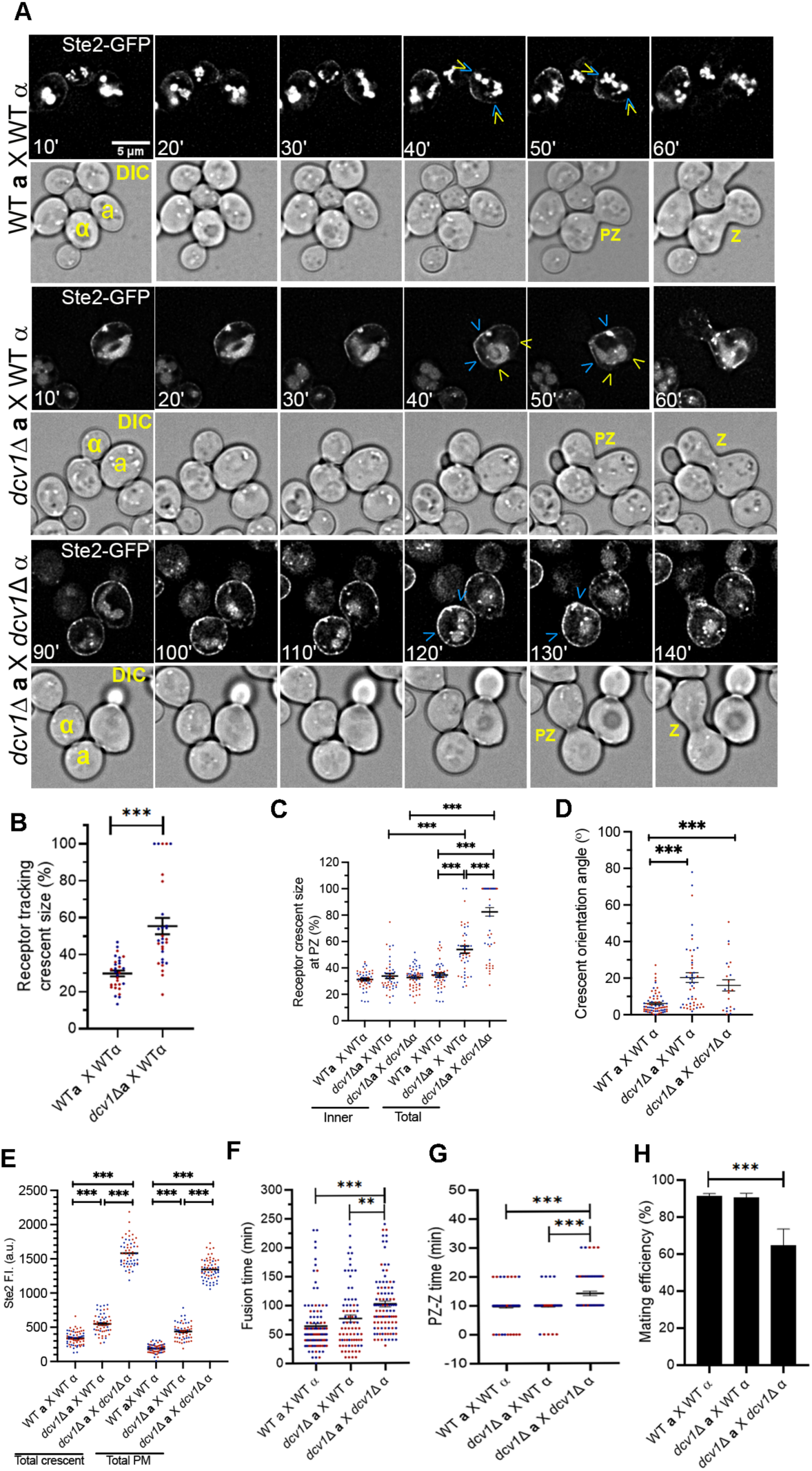
Mating *dcv1*Δ cells exhibit defects in receptor polarization, receptor orientation, and cell fusion. Log-phase *MAT***a** *DCV1* and *MAT***a** *dcv1Δ* cells expressing Ste2-GFP were mixed with an equal number of *MAT*α cells marked with Bud1-RFP and imaged until fusion. (A) The receptor is highly depolarized in mating *dcv1*Δ cells. Center slices of representative images are shown. The mating partners are labeled **a** and α in the DIC images; PZ, prezygote; Z, zygote. Blue arrowheads indicate the edges of inner crescents and yellow arrowheads indicate the edges of total crescents. (B) Tracking receptor crescents on the PM of mating *MAT***a** *dcv1*Δ cells are depolarized. Total crescent size was measured at midpoint between the DS and CS. Data points represent the crescent sizes measured from two independent experiments, indicated by color. Horizontal lines and error bars indicate mean ± s.e.m. (control = 30.8 ± 1.6, unilateral = 55.4 ± 4.4. n = 30. \*\*\**p* < 0.0001). (C) Receptor crescents on the PM of mating *MAT***a** *dcv1*Δ cells at the PZ stage are depolarized and lack clear boundaries. The inner crescent measurement includes only the well-demarcated region of highest receptor concentration; the total crescent measurement includes the inner crescent plus detectable receptor signal that extends beyond it. Total crescent size was measured only if inner crescents lacked a clear boundary. Data points represent the crescent sizes measured from two independent experiments, indicated by color. Horizontal lines and error bars indicate mean ± s.e.m. (Inner: control = 31.3 ± 1.1, unilateral = 33.9 ± 1.9, bilateral = 32.8 ± 1; total: control = 34.6 ± 10.7, unilateral = 53.9 ± 2.8, bilateral = 82.4 ± 3.2. n = 45. \*\*\**p* < 0.0001). (D) *dcv1*Δ confers a defect in receptor crescent alignment with the mating partner. Crescent orientation is defined as the angle between the center of the fusion zone and the center of the total receptor crescent at the PZ stage. Data points represent crescent orientation angles measured in two independent experiments, indicated by color. Horizontal lines and error bars indicate mean ± s.e.m. (control = 6.1 ± 0.7; unilateral = 20.3 ± 2.7; bilateral = 16 ± 3.08. n: control = 68; unilateral = 48; bilateral = 22. \*\*\**p* < 0.0001). (E) Mating *dcv1*Δ cells express aberrantly high levels of Ste2-GFP on the PM. Data points represent the mean fluorescent intensities in the polarized receptor crescent and in the total PM measured in two independent experiments, indicated by color. Horizontal lines and error bars indicate mean ± s.e.m. (Total crescent: control = 339.7 ± 11.9, unilateral = 548 ± 19.9, bilateral = 1582 ± 26.3; total PM: control = 190.4 ± 7.3, unilateral = 439 ± 16.8, bilateral = 1345 ± 19.9. n *≥* 53. \*\*\**p* < 0.0001). (F) *dcv1*Δ bilateral mating mixtures exhibit a delay in zygote formation. Data points represent the time in minutes from the start of imaging to fusion measured in two independent experiments, indicated by color. Horizontal lines and error bars represent mean ± s.e.m. (control = 64 ± 4.5; unilateral = 77.1 ± 6.2; bilateral = 101.9 ± 5.1. (n *≥* 95. \*\*\**p* < 0.0001; \*\**p* = 0.002). (G) Cells in *dcv1*Δ bilateral mating mixtures exhibit a delay in prezygote to zygote (PZ to Z) progression. Data points represents the time in minutes taken to progress from PZ to Z measured in two independent experiments, indicated by color. Lines and error bars represent mean ± s.e.m. (control = 9.8 ± 0.5; unilateral = 10 ± 0.6; bilateral = 14.14 ± 0.7. n *≥* 78. \*\*\**p* < 0.0001) (H) Mating efficiency is significantly reduced in *dcv1*Δ bilateral mating mixtures. Mating efficiency was calculated as a percentage of observed zygotes/potential zygotes (potential partners within 3 µm of each other). Bar graphs represents the mean mating efficiency ± s.e.m. measured in two independent experiments. (control = 91.6 ± 1.1; unilateral = 90.7 ± 2.1; bilateral = 64.9 ± 8.7. n, expected zygotes: control = 101; unilateral = 91; bilateral = 110; \*\*\**p*_[chi sq]_ < 0.0001).

Mutations that confer mating phenotypes are often more penetrant and/or expressive when both mating partners are mutant (Erdman et al., 1998; Gehrung and Snyder, 1990; Kurihara et al., 1994). Indeed, the receptor localized over the entire PM in about two thirds of the *MAT***a** *dcv1*Δ cells mating with *MATα dcv1*Δ cells, with Ste2-GFP forming enriched inner crescents in about 94% of the cells examined (Fig. 4A,C). In the remaining third, the majority (60%) of cells failed to center the total receptor crescents around the eventual fusion site (Fig. 4D). Moreover, *MAT***a** *dcv1*Δ cells that formed zygotes in bilateral mating mixtures displayed considerably higher PM Ste2-GFP signals than the elevated signal seen in *MAT***a** *dcv1*Δ cells mating with WT partners (Fig. 4E). We also found that the time to cell fusion in the bilateral mating mixtures was significantly greater than in the control and unilateral mixtures (Fig. 4F), consistent with the observation that zygote formation is delayed when *MAT***a** cells are unable to polarize the receptor (Ismael et al., 2016). Notably, the *MAT***a** *dcv1*Δ/*MATα dcv1*Δ prezygotes were significantly delayed in progressing to zygotes (Fig. 4G), suggesting a defect in localized cell wall degradation and/or PM fusion. This observation is consistent with the proposed role for an interaction between the activated receptors of mating partners in these processes (Shi et al., 2007). An additional indication that Dcv1 plays an important role in mating was the significantly reduced percentage of zygotes formed in the bilateral assays (Fig. 4H). Inspection of the time-lapse images revealed that, unlike *MAT***a** cells in the control or unilateral assays, *MAT***a** *dcv1*Δ and *MATα dcv1*Δ cells rarely mated unless they were initially in contact with a potential partner.

### Dcv1 contributes to the localization of the endocytic adaptor, Sla1

Directed delivery, slow diffusion, and localized endocytosis of PM proteins act together to establish and maintain polarity sites in yeast (Valdez-Taubas and Pelham, 2003). The internalization of Ste2 through receptor-mediated endocytosis (RME) plays a crucial role in its polarization during the chemotropic response (Suchkov et al., 2010). Sla1 is the primary adaptor for Ste2 RME (Goode et al., 2015; Howard et al., 2002) and thereby serves as a marker of receptor endocytosis. *dcv1*Δ cells exhibit a loss of receptor polarity characterized by larger receptor crescents that do not have clearly demarcated end points. To determine whether this phenotype is associated with a defect in RME, we took time-lapse images of mating *MAT***a** *DCV1* (WT) and *MAT***a** *dcv1Δ* cells engineered to co-express Ste2-GFP and Sla1-RFP (Fig. 5A). Localization of the reporters was scored at the PZ stage. Sla1 puncta were classified as either flanking the fusion zone (FZ localization), outside the FZ but within the mating projection as defined by the ends of the Ste2-GFP crescent (apical localization), or outside the mating projection (ectopic localization). Consistent with previous results (Wang et al., 2019), over 90% of the WT cells polarized Sla1 to the mating projection, with most showing exclusive localization to the FZ. In contrast, less than half as many *dcv1*Δ cells exhibited Sla1 puncta exclusively at the FZ, whereas the proportions of mutant cells showing apical and ectopic Sla1 localization were significantly increased (Fig. 5B). Notably, mislocalization of Sla1 in mating *dcv1*Δ cells did not correlate with either an increase in mean receptor crescent size (Fig. 5C) or crescent misorientation (Fig. 5D). These data suggest that the receptor polarity phenotypes observed in *dcv1*Δ cells are not due to a defect in RME. We infer that the absence of Dcv1 independently affects receptor and Sla1 localization.

**Figure 5:**
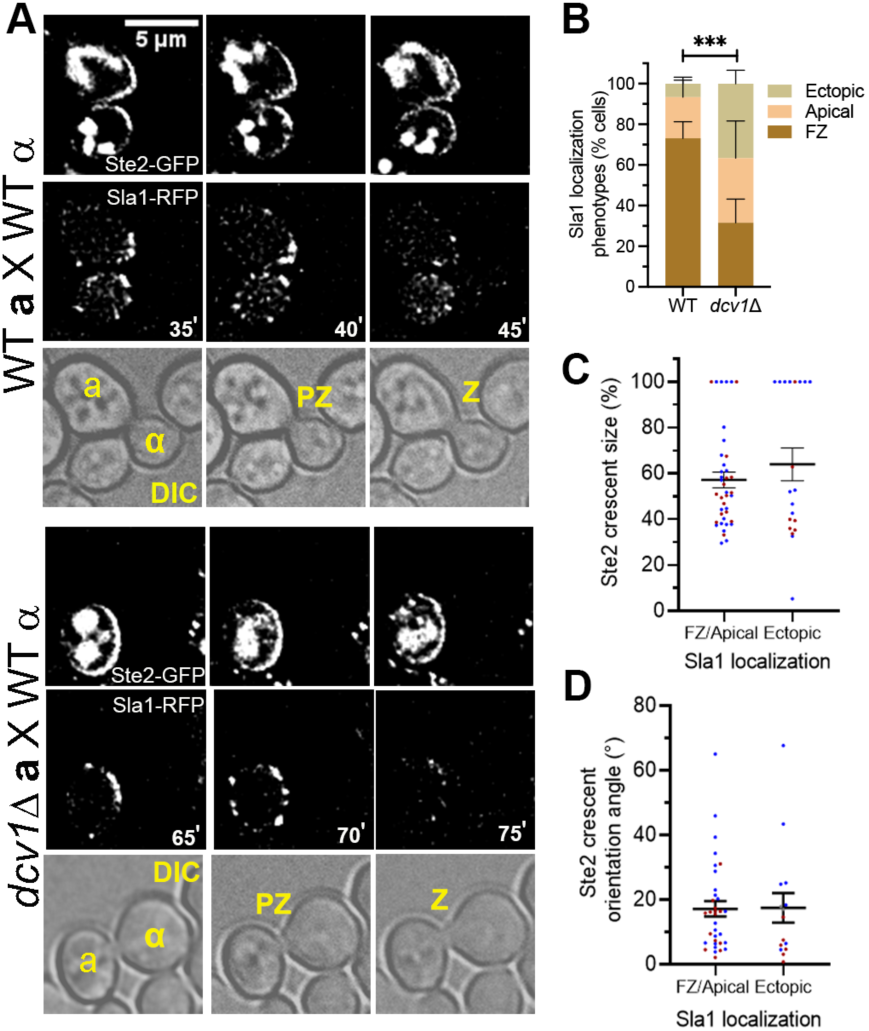
*dcv1*Δ causes mislocalization of Sla1 in mating cells independent of its effects on receptor polarization and orientation. Log-phase *MAT***a** *DCV1* and *MAT***a** *dcv1Δ* cells co-expressing Ste2-GFP and Sla1-RFP were mixed with an equal number of *MAT*α cells and imaged every five minutes until fusion. (A) Sla1 is mislocalized in *dcv1*Δ prezygotes. Center slices of representative images are shown. The mating partners are labeled **a** and α in the DIC images; PZ, prezygote; Z, zygote. (B) Sla1 localization outside of the fusion zone is dramatically increased in *dcv1*Δ prezygotes. The Sla1 localization phenotypes were scored at the PZ stage as follows: all puncta within the fusion zone (FZ, light brown); all puncta within the front 40% (Apical, salmon); some puncta outside the apical region (Ectopic, brown). Bar graphs represent the mean percentage of cells ± s.e.m. in two independent experiments (WT: FZ = 73.1 ± 8.1, apical = 20.2 ± 9.8, ectopic = 6.7 ± 1.7; *dcv1*Δ: FZ = 31.6 ± 11.6, apical = 31.8 ± 18.2, ectopic = 36.6 ± 6.6. n ≥ 62. \*\*\**p* < 0.0001). (C-D) Ste2-GFP crescent size and orientation were determined in *dcv1*Δ prezygotes and Sla1 localization categorized as either FZ/apical or ectopic. (C) Receptor depolarization in *dcv1*Δ prezygotes does not correlate with ectopic Sla1 localization. Data points represent crescent sizes for each category measured in two independent experiments, indicated by color. Horizontal lines and error bars represent mean ± SEM (FZ/apical = 57.1 ± 3.4; ectopic = 63.9 ± 7.2. n: FZ/apical = 40; ectopic = 20. *p* = 0.33). (D) Receptor crescent misalignment in *dcv1*Δ prezygotes does not correlate with ectopic Sla1 localization. Data points represent crescent orientation angle for each category measured in two independent experiments, indicated by color. Horizontal lines and error bars represent mean ± SEM (FZ /apical = 17.1 ± 2.4; ectopic = 17.4 ± 4.6. n: FZ/apical = 34; ectopic = 15. *p* = 0.95).

### Dcv1 contributes to the localization of the polarisome component, Spa2, in mating cells

A multiprotein complex known as the polarisome (Pruyne and Bretscher, 2000) must be appropriately positioned at the chemotropic site (CS) to localize the secretory vesicles for proper shmoo formation and cell fusion (Bidlingmaier and Snyder, 2004). Pheromone-treated cells lacking the polarisome scaffold protein Spa2 form broad mating projections (Banavar et al., 2018; Gehrung and Snyder, 1990), a phenotype also exhibited by *dcv1*Δ cells. To ask whether the effect of *dcv1*Δ on shmoo morphology involves the polarisome, we assayed polarisome localization in time-lapse images of mating cells using Spa2-GFP as a proxy. In both *MAT***a** *DCV1* and *MAT***a** *dcv1Δ* cells, Spa2-GFP localized as a single spot to the neck of cytokinetic cells (Fig. 6A), as expected from previous reports (Dobbelaere and Barral, 2004). Whereas most mating WT cells polarized Spa2-GFP as a single spot at all time points, the majority of *dcv1*Δ cells exhibited more than one Spa2-GFP spot for at least one time point (Fig. 6A,B). The single Spa2-GFP spot in WT cells either disappeared from the neck and reappeared at the CS, which we call jumping, or moved persistently from the neck to the CS, which we call tracking. In a small fraction of WT cells, Spa2-GFP did not move persistently toward the CS; rather, it vacillated prior to its stabilization (Fig. 6A,C). We call this behavior wandering. A higher incidence of Spa2-GFP jumping and wandering was observed in the mutant cells, with a corresponding decrease in tracking (Fig. 6C). *dcv1*Δ cells also took significantly longer than control cells to progress from the neck localization of Spa2-GFP to cell fusion. Consistent with a higher incidence of Spa2-GFP wandering, most of this delay occurred during Spa2-GFP movement from the neck to the CS (Fig. 6D).

**Figure 6:**
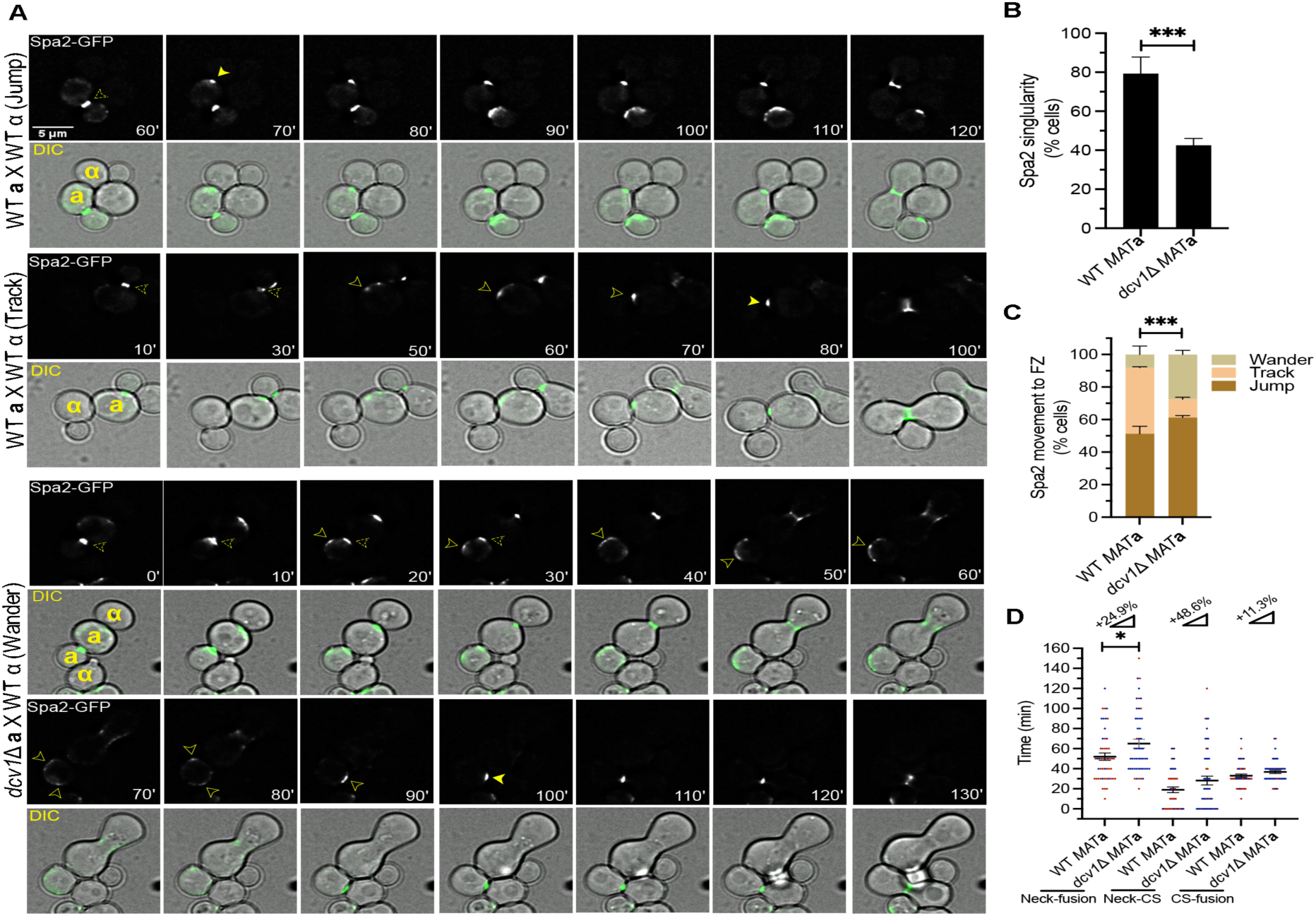
*dcv1*Δ confers Spa2 localization defects. Log-phase *MAT***a** *DCV1* and *MAT***a** *dcv1Δ* cells expressing Ste2-GFP were mixed with an equal number of *MAT*α cells expressing Bud1-RFP and imaged every 10 minutes from cytokinesis to fusion. (A) Spa2 is mislocalized in mating *dcv1Δ* cells. Center slices of representative images are shown. The mating partners are labeled **a** and α in the DIC images. Dashed arrowheads indicate Spa2 localization at the neck; solid arrowheads indicate mobile (tracking/wandering) Spa2; filled arrowhead indicate stable Spa2 localization at the CS. (B) *dcv1*Δ cells are defective in polarizing Spa2 to a single site. Cells were characterized as having either one or more-than-one polarized Spa2 patch at each time point. Bar graphs represent the mean percentage of cells with a single polarized Spa2 patch ± s.e.m. measured in two independent experiments (WT = 79.31 ± 5.98; *dcv1*Δ = 42.5 ± 2.5. n = 49. \*\*\**p*_[chi sq]_ = 0.0002.). (C) Spa2 wandering is more likely to occur in *dcv1*Δ cells. Spa2 movement from the neck to the CS was characterized as either jumping (brown), tracking (persistent movement toward CS, salmon) or wandering (movement away from the CS for at least one time point, light brown). Bar graphs represent the mean percentage of cells in each category ± s.e.m. measured in two independent experiments (WT: jump = 51.27 ± 4.61, track = 40.59 ± 0.59, wander = 8.14 ± 5.2; *dcv1*Δ: jump = 61.25 ± 1.25, track = 11.25 ± 1.25, wander = 27.5 ± 2.5. n = 49; \*\*\**p*_[chi sq]_ < 0.0001.). (D) *dcv1*Δ confers a delay in zygote formation that correlates with an increase in the time it takes Spa2 to move from the neck to the CS. Data points represent the following time intervals in minutes: Spa2 translocation from the neck to cell fusion (neck-fusion), Spa2 translocation from the neck to the CS (neck-CS), and Spa2 stabilization at the CS to cell fusion (CS-fusion). Measurements are from two independent experiments, indicated by color. Ramp symbols and numbers represent the percent increase in the respective time intervals observed in *dcv1*Δ cells. Lines and error bars represent mean ± s.e.m. (WT: neck-fusion = 46.7 ± 3, neck-CS = 13.7 ± 2.3, CS-fusion = 33.1 ± 1.7; *dcv1*Δ: neck-fusion = 56.6 ± 4.1, neck-CS = 19.8 ± 4, CS-fusion = 36.8 ± 1.6. n = 49. \**p* = 0.027).

### Dcv1 contributes to the localization of the fusion protein, Fus1, in mating cells

Mating cells generate a tightly focused FZ at the tip of the mating projection, where they ultimately contact their partners. The FZ is highly enriched in fusion-specific proteins (e.g., Fus1, Fus2, Fig1, and Prm1), and the PM lipids ergosterol and sphingolipids (Merlini et al., 2013). Fus1 functions as a scaffolding protein: it assembles other fusion-specific proteins and regulates polarized secretion of the enzymes that locally degrade the PM and cell wall (Bagnat and Simons, 2002; Nelson et al., 2004; Trueheart et al., 1987). As shown in Figure 6D, *dcv1*Δ conferred an increase in the time between cytokinesis and cell fusion. This could reflect a delay in localizing the fusion machinery to the CS, or a defect in the function of the fusion machinery at the CS, or both. To distinguish these possibilities, we imaged Fus1-GFP in mating cells as a proxy for the fusion machinery. In most *MAT***a** *DCV1* cells, Fus1-GFP initially formed a polarized crescent at the FZ, which then condensed into a bright spot (hereafter, the cap) about eight minutes before fusion (Fig. 7A,B). In a small fraction of WT cells (13%), the polarized Fus1-GFP crescent formed away from the FZ (ectopically), then either jumped or tracked to the FZ prior to cap formation (7C). A significantly higher fraction of the *dcv1*Δ cells polarized Fus1-GFP ectopically (37%), and although all such Fus2-GFP crescents eventually localized to the FZ in advance of cap formation, we saw a high incidence of reporter wandering at the expense of tracking (Fig. 7A-C), similar to the behavior of Spa2-GFP in mutant cells (Fig. 6A,C). Moreover, *dcv1*Δ cells took significantly longer than control cells to progress from emergence of the Fus1 crescent to fusion. The higher incidence of ectopic Fus1-GFP localization, followed by wandering in the mutant cells, is consistent with the delay occurring entirely before cap formation. The cap-to-fusion interval was indistinguishable in control and mutant cells (Fig. 7D). Notably, the delay in progression of Fus1 crescent formation to cell fusion was comparable to the delay in progression of Spa2 neck localization to cell fusion. These data suggested that the mating of *dcv1*Δ cells is delayed due to their difficulty in localizing the polarizome and fusion machinery to the CS.

**Figure 7:**
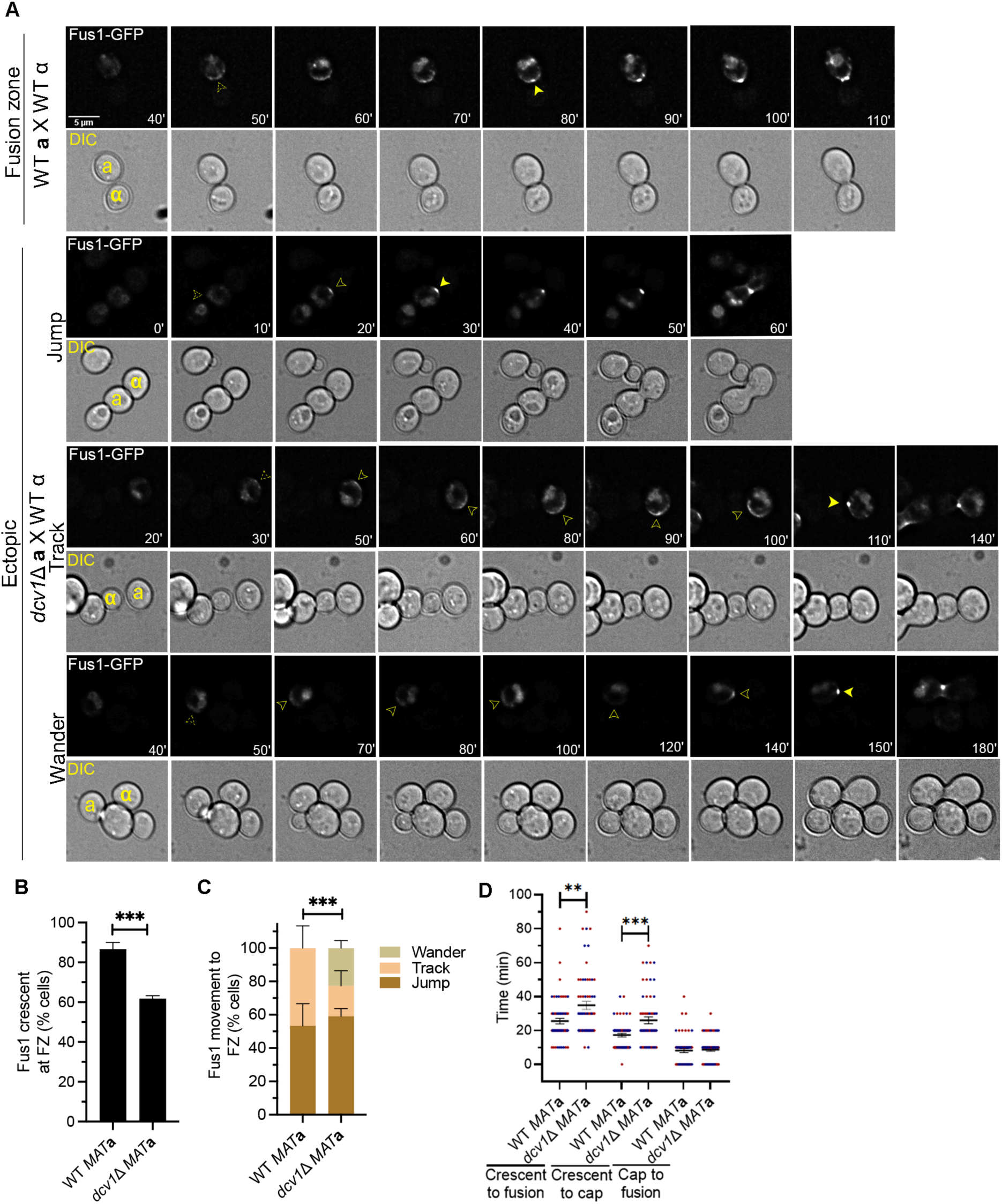
*dcv1*Δ confers defects in Fus1 localization. Log-phase *MAT***a** *DCV1* and *MAT***a** *dcv1Δ* cells expressing Fus1-GFP were mixed with an equal number of *MAT*α cells expressing Bud1-RFP and imaged every 10 minutes from cytokinesis to fusion. (A) Fus1 is mislocalized in mating *dcv1*Δ cells. Center slices of representative images are shown. The mating partners are labeled **a** and α in the DIC images. Dashed arrowheads indicate Fus1-GFP crescent emergence, solid arrowheads indicate mobile (tracking/wandering) Fus1-GFP, and filled arrowheads indicate Fus1-GFP cap emergence at the FZ. (B) The incidence of Fus1-GFP crescent emergence at the FZ is significantly reduced in *dcv1*Δ cells. Cells were scored according to whether the Fus1-GFP crescent was first visible at the FZ or away from it (ectopic). Bar graphs represent the mean percentage of cells showing initial polarization of Fus1-GFP at the FZ ± s.e.m., measured in two independent experiments (WT = 86.67 ± 3.33; *dcv1*Δ = 61.83 ± 1.5. n = 60. \*\*\**p*_[chi sq]_ = 0.0002.). (C) *dcv1*Δ confers a defect in Fus1-GFP tracking. Ectopic Fus1-GFP movement to the FZ was characterized as either jumping (disappearance from the ectopic site and reappearance at the FZ, brown), tracking (persistent movement toward FZ, salmon), or wandering (backtracking for at least one time point prior to stable FZ localization, light brown). Bar graphs represent the mean percentage of cells in each category ± s.e.m., measured in two independent experiments (WT: jump = 53.3 ± 13.3, track = 46.7 ± 13.3; *dcv1*Δ: jump = 59.1 ± 4.5, track = 18.2 ± 9.1, wander = 22.7 ± 4.6. n: WT = 8; *dcv1*Δ = 22. \*\*\**p*_[chi sq]_ = 0.0001). (D) Fus1-GFP cap emergence is significantly delayed in *dcv1*Δ cells. Total time taken from Fus1-GFP crescent emergence to cell fusion, Fus1-GFP crescent emergence to cap emergence, and Fus1-GFP cap emergence to cell fusion were determined. Data points represent the time taken in minutes for each category, measured in two independent experiments, indicated by color. Lines and error bars represent mean ± s.e.m. (WT: crescent to fusion = 25.5 ± 1.6, crescent to cap = 17.3 ± 1, cap to fusion = 8.2 ± 1.1; *dcv1*Δ: crescent to fusion = 34.8 ± 2.4, crescent to cap = 26 ± 2, cap to fusion = 8.8 ± 1. n = 60. \*\**p* = 0.002; \*\*\**p* = 0.0002).

### Blocking ergosterol synthesis partially phenocopies the effects of *dcv1*Δ in mating cells

Polarized lipids play important roles in organizing signaling and polarity proteins on the PM during mating. The primary PM lipid, ergosterol, is critical for mating functions such as localization of the Ste5 signaling scaffold protein to the tip of mating projection, polarized growth, and cell fusion (Jin et al., 2008). Ergosterol biosynthetic mutants, such as *erg6*Δ cells, exhibit the following phenotypes: 1) rounded shmoo morphology; 2) defects in polarizing sterols and Fus1 to the mating projection and FZ; 3) fusion delays (Aguilar et al., 2010; Bagnat and Simons, 2002; Jin et al., 2008; Tiedje et al., 2007). Because *dcv1*Δ cells exhibit similar phenotypes, we wondered whether blocking ergosterol synthesis would affect receptor polarization. To answer this question, we measured receptor crescent sizes and orientation angles in mating *MAT***a** *erg6*Δ cells expressing Ste2-GFP (Fig. 8A-C). Like *dcv1*Δ, *erg6*Δ conferred significant, although less severe, defects in receptor polarization and orientation. *erg6*Δ cells also formed sharply defined crescents within larger crescents that lacked clear boundaries, like *dcv1*Δ cells; surprisingly, the mean inner-crescent size in *erg6*Δ cells was significantly smaller than the crescents formed by WT cells (Fig. 8B). Unlike *dcv1*Δ cells, *erg6*Δ cells displayed normal levels of Ste2-GFP on the PM (Fig. 8D). In contrast to both WT *MAT***a**/*MAT*α and *MAT***a** *dcv1*Δ/*MAT*α *DCV1* zygotes, the majority of *MAT***a** *erg6*Δ/*MAT*α *ERG6* zygotes budded away from the FZ (Fig. 8E). Taken together, these data demonstrate that *erg6*Δ partially phenocopies *dcv1*Δ, suggesting that the polarized mating function defects observed in *dcv1*Δ cells can be at least partially attributed to altered PM lipid composition.

**Figure 8:**
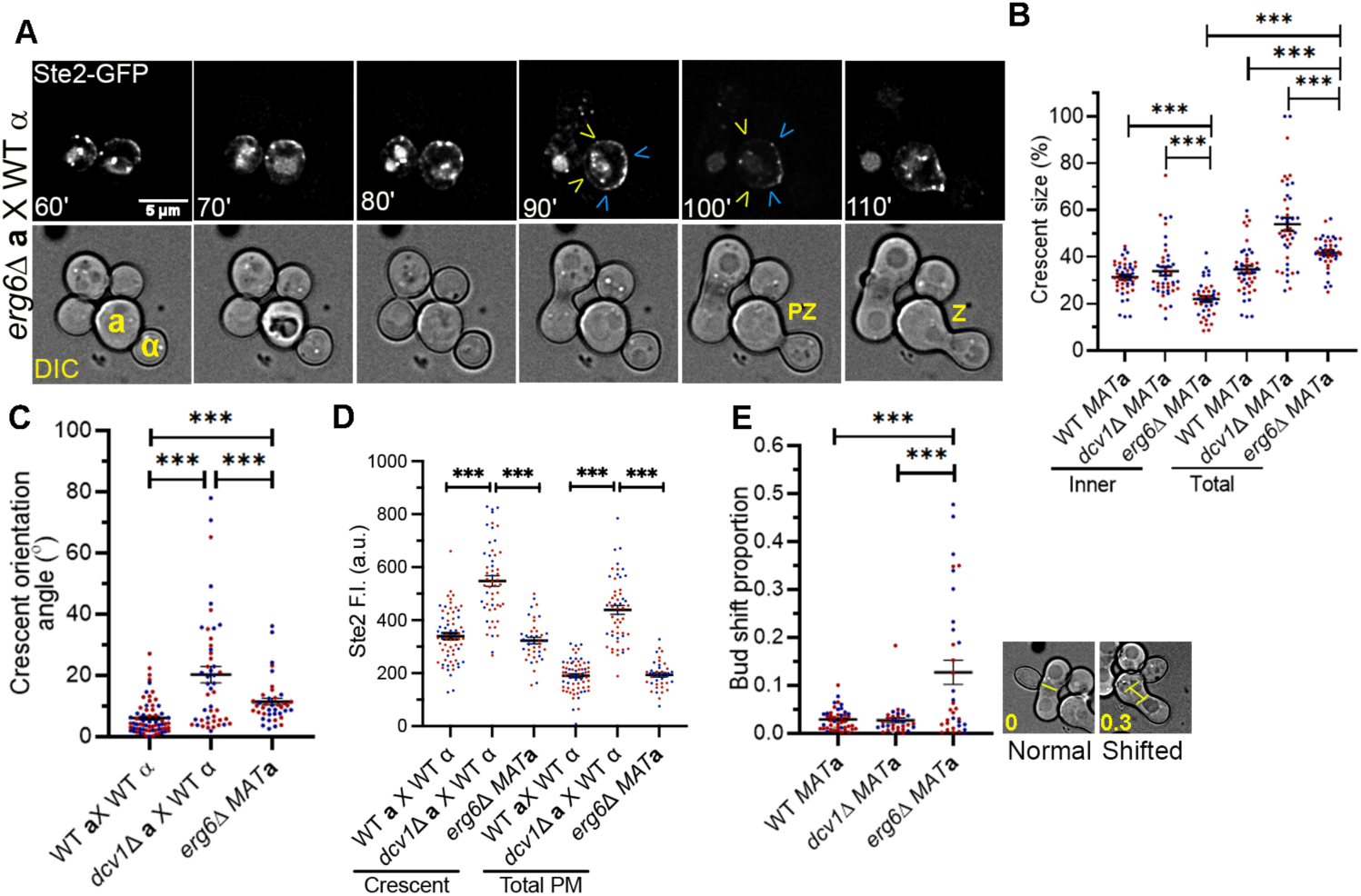
*erg6*Δ partially phenocopies *dcv1*Δ. Log-phase *MAT***a** *erg6*Δ cells expressing Ste2-GFP were mixed with an equal number of *ERG6* (WT) *MAT*α cells expressing Bud1-RFP and imaged every ten minutes until fusion. (A-B) The receptor is depolarized in mating *erg6*Δ cells. (A) Center slices of representative images are shown. The mating partners are labeled **a** and α in the DIC images; PZ, prezygote; Z, zygote. Blue arrowheads indicate the edges of the inner crescents. Yellow arrowheads indicate the edges of the total crescents. (B) The polarized receptor crescents on the PM of mating *MAT***a** *erg6*Δ cells lack clear boundaries. Data points represent the crescent sizes measured in two independent experiments, indicated by color. Lines and error bars represent mean ± s.e.m. (Inner: WT = 31.3 ± 1, *dcv1*Δ = 33.9 ± 1.9, *erg6*Δ = 21.9 ± 1.2; total: WT = 34.6 ± 1.6, *dcv1*Δ = 53.9 ± 2.8, *erg6*Δ = 41.7 ± 1.1. n *≥* 40. \*\*\**p* ≤ 0.0006). Data for the WT and unilateral mutant crosses duplicated from Fig. 4C for convenience. (C) *erg6*Δ confers a defect in receptor-crescent alignment with the mating partner. Crescent orientation angle was measured as described in Fig. 4D. Data points represent crescent orientation angles measured in two independent experiments, indicated by color. Horizontal lines and error bars indicate mean ± s.e.m. (WT = 6.1 ± 0.7; *dcv1*Δ = 20.3 ± 2.7; *erg6*Δ = 11.4 ± 1.2. n *≥* 40. ***p < 0.0001). Data for the WT and unilateral mutant crosses duplicated from Fig. 4D for convenience. (D) Mating *erg6*Δ cells express normal levels of Ste2-GFP on the PM. Data points represent the mean fluorescent intensities (F.I.) in the polarized receptor crescent and in the total PM measured in two independent experiments, indicated by color. Horizontal lines and error bar indicate mean ± s.e.m. (Crescent: WT = 339.7 ± 11.9, *dcv1*Δ = 548 ± 19.9, *erg6*Δ = 323.1 ± 12.4; total PM: WT = 190.4 ± 7.4, *dcv1*Δ = 439 ± 16.8, *erg6*Δ = 193.7 ± 8.3. n *≥* 40. ***p < 0.0001). Data for the WT and unilateral mutant crosses duplicated from Fig. 4E for convenience. (E) Zygotic buds are rarely positioned at the FZ in *erg6*Δ cells. Images are representative of a bud positioned at the FZ (left) and a shifted bud (right). The bud-shift proportions (yellow numbers on the DIC images) were obtained by dividing the distance between the center of the FZ and the center of the bud (yellow lines) by the zygote length. Data points represent the bud-shift proportions measured in two independent experiments, indicated by color. Horizontal lines and error bars indicate mean bud-shift proportion ± s.e.m. (WT = 0.03 ± 0.003; *dcv1*Δ = 0.03 ± 0.006; *erg6*Δ = 0.13 ± 0.025. n *≥* 32. ***p ≤ 0.0003).

## DISCUSSION

The establishment of front-rear polarity is integral to directed cell migration and growth. How this polarity is generated and maintained is not fully understood. In this study, we identified a novel player in cell polarization during yeast mating. Deletion of *DCV1*, a claudin homolog, adversely affected multiple cellular processes: protein and actin-cable polarization, PM lipid enrichment, cell integrity, shmoo morphology, gradient tracking, and fusion with mating partners.

Our results implicate Dcv1 in organizing and/or maintaining front-rear polarity in mating cells. The polarized receptor crescents, which are typically restricted to the front one-third of the cell, extended into the rear domain, and often spanned the entire cell periphery in mating *dcv1*Δ cells. Notably, the receptor crescents lacked a sharp boundary: although the inner crescent size was comparable to control cells, the outer crescent gradually decreased along the PM to a level below our detection. Similarly, in pheromone-treated cells, the enrichment of PIP2 and PS extended beyond the front domain, and we observed ectopic localization of PM ergosterol and the RME marker, Sla1. These results are consistent with the idea that Dcv1 provides a segregation or barrier function, without which the front and rear domains of pheromone-stimulated cells are not clearly demarcated.

When and how are the front-rear membrane domains established in pheromone-stimulated yeast? We previously reported that mating cells assemble a gradient tracking machine (GTM) consisting of sensory, polarity, and secretory proteins at the DS, and that the GTM then redistributes, or tracks, to the CS (Wang et al., 2019). Tracking is driven by directed vesicle delivery on the upgradient side of the GTM coupled with endocytosis on its downgradient side. Here we found that in cells responding to isotropic pheromone, PS and PIP2 translocated from the neck to the DS, the site of GTM assembly, prior to morphogenesis. This observation raises the possibility that the GTM consists of lipids as well as proteins, and that both are necessary for chemosensing and cell polarization. Consistent with this conjecture, it has been demonstrated that specific lipids are required for the polarization/retention of signaling and polarity proteins: Sterols are required for the formation of membrane domains (Bagnat and Simons, 2002) that promote shmoo-tip localization of the MAPK scaffold, Ste5 (Jin et al., 2008), and the p21- activated kinase, Ste20 (Tiedje et al., 2007); PS polarization facilitates proper shmoo-tip clustering of Cdc42 (Fairn et al., 2011; Sartorel et al., 2018; Yeung et al., 2008); and PIP2 is required for efficient targeting and/or organization of various proteins involved in polarized growth (e.g., Ste20, the Cdc42 effector Gic2, septins, and the exocyst component Exo70 (He *et al*., 2007a; Takahashi and Pryciak, 2007; Orlando *et al*., 2008; Bertin *et al*., 2010). We speculate that the GTM is a lipid-protein domain that establishes front-rear polarity when it is assembled at the DS of cells treated with isotropic pheromone and when it stabilizes at the CS of gradient-stimulated mating cells.

How does Dcv1 contribute to front-rear polarity? Although we cannot yet answer this question in detail, our data and the results published by other groups suggest that the primary defect in *dcv1*Δ cells, which could plausibly result in all the phenotypes we observed, is the mislocalization of the PM lipids. The alternative explanation – that *dcv1*Δ independently affects multiple proteins and pathways – is much less economical. We considered the possibility that the attenuation of receptor polarity in *dcv1*Δ cells was due to mislocalization of Sla1, or vice versa, but found no correlation between the “trailing receptor” and ectopic Sla1 phenotypes. The unrelated occurrence of these phenotypes argues against an interdependent mechanism, and therefore, for a common cause that generates independent effects. Additional genetic evidence associating Dcv1 with PM lipids includes our finding that *erg6*Δ, which blocks ergosterol biosynthesis, phenocopied many of the effects of *dcv1*Δ in mating cells, and the severe synthetic growth defects that result when *dcv1*Δ is combined with deletion of *IRS4* (which regulates PIP2 levels) or *CYB5* (an electron donor in sterol and lipid biosynthesis) (Costanzo et al., 2016).

In principle, Dcv1 could influence PM domains by affecting the distribution, metabolism, exchange, and/or flipping of lipids. However, membrane domain formation in eukaryotes is thought to largely depend on protein scaffolding, wherein homomeric and heteromeric protein complexes directly bind specific lipids, and protein fencing, wherein heteromeric protein complexes prevent lateral diffusion of specific proteins and lipids between PM domains. Scaffolding is exemplified by Pil1 and Lsp1, which self-assemble into a complex that preferentially binds phosphoinositides to generate the eisosomes (Olivera-Couto and Aguilar, 2012; Walther et al., 2006). The roles of claudins in tight junctions between epithelial cells and the septin ring that separates mother and daughter yeast cells are examples of fencing. The localization of Dcv1 away from the receptor as pheromone-treated cells begin morphogenesis is consistent with either of these paradigms: Dcv1 could act as a rear-domain scaffold, binding one or more specific lipids; or, by virtue of being a four-pass integral membrane protein, it could serve as a fence, preventing lateral diffusion of the receptor and other GTM components beyond the mating projection. It is also possible that Dcv1 directly affects PM lipid composition. The <u>o</u>xy<u>s</u>terol binding protein <u>h</u>omologue (OSH) proteins, which are involved in organizing and trafficking sterols on the membrane (Georgiev et al., 2011), bind and exchange PM sterols for PI4P. Of note, a region of Dcv1 spanning its predicted second and third intracellular loops (residues 112 – 172) is 32% identical and 51% similar to the conserved oxysterol binding domain (residues 641 – 706) of the yeast Osh3 protein, which promotes PI4P polarity (Omnus et al., 2020). Dni1, Dni2, and Sur7 are examples of claudin-like fungal proteins that have been implicated in PM organization and cell wall remodeling (Alvarez et al., 2008; Clemente-Ramos et al., 2009; Curto et al., 2018).

The segregation of Dcv1 and the receptor as cells begin to shmoo, together with the loss of distinct receptor boundaries in *dcv1*Δ cells, strongly suggests that Dcv1 plays a key role in maintaining pheromone-induced front-rear polarity. It is less clear, however, whether Dcv1 contributes to the establishment of the front and rear domains. Although it remains to be determined, we favor the idea that front determination begins with assembly of the GTM – whose components include PIP2, PS, and possibly other lipids – at the DS. In cells treated with isotropic pheromone, the GTM remains at the DS, where it triggers shmooing (front domain formation); in mating cells, the GTM aligns with the gradient source before triggering robust polarized secretion and the consequent maturation of front-rear polarity. Two pieces of evidence suggest that Dcv1 does indeed play a role in GTM function. First, the receptor was depolarized and lacked sharp boundaries while tracking to the CS in *dcv1*Δ cells. Second, the Spa2 polarisome protein exhibited significant tracking defects (wandering, multiple foci) in *dcv1*Δ cells. Thus, the novel rear-domain protein and claudin homolog, Dcv1, may be important for both the establishment and maintenance of front-rear polarity.

## ACKNOWLEDGEMENTS

We thank Sergio Grinstein, Elizabeth Grayhack, and Robert Arkowitz for providing plasmids. We thank members of the Stone lab for helpful discussions and critical reading of the manuscript. This work was supported by National Science Foundation grants 1415589 and 1818067 (to D.E.S.). The authors declare no competing financial interests.

---

**Figure S1:**
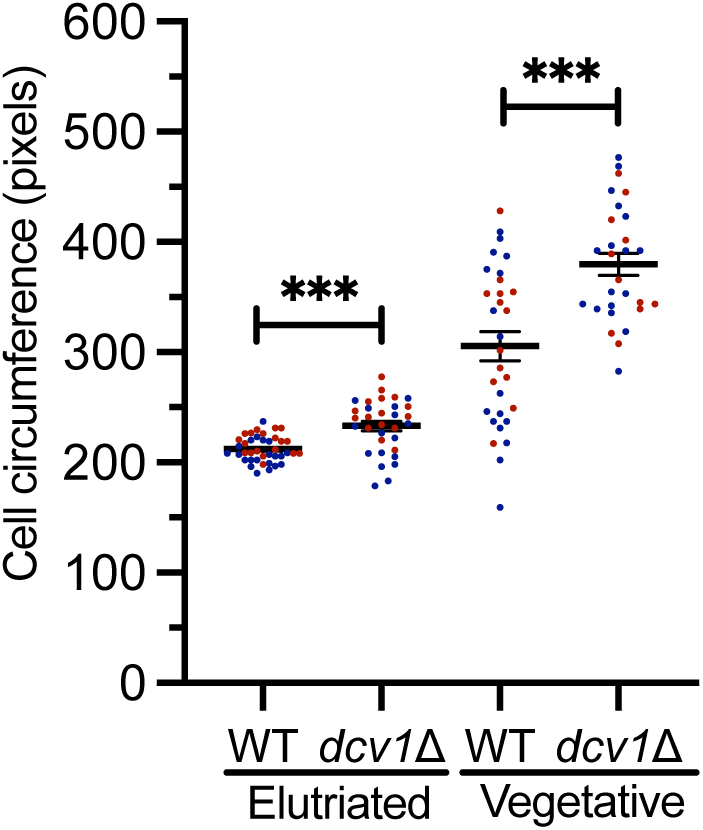
*dcv1*Δ confers an increase in cell size. The cell circumferences of elutriated and log-phase vegetative (mother and single) cells were measured using ImageJ. Data points represent the cell circumference in pixels measured in two independent experiments, indicated by color. Lines and error bars represent mean ± s.e.m. (WT: elutriated = 212.1 ± 1.8, vegetative = 305.6 ± 13.1; *dcv1*Δ: elutriated = 233.2 ± 4.3, vegetative = 379.6 ± 9.9. n *≥* 28. \*\*\**p* < 0.0001).

**Figure S2:**
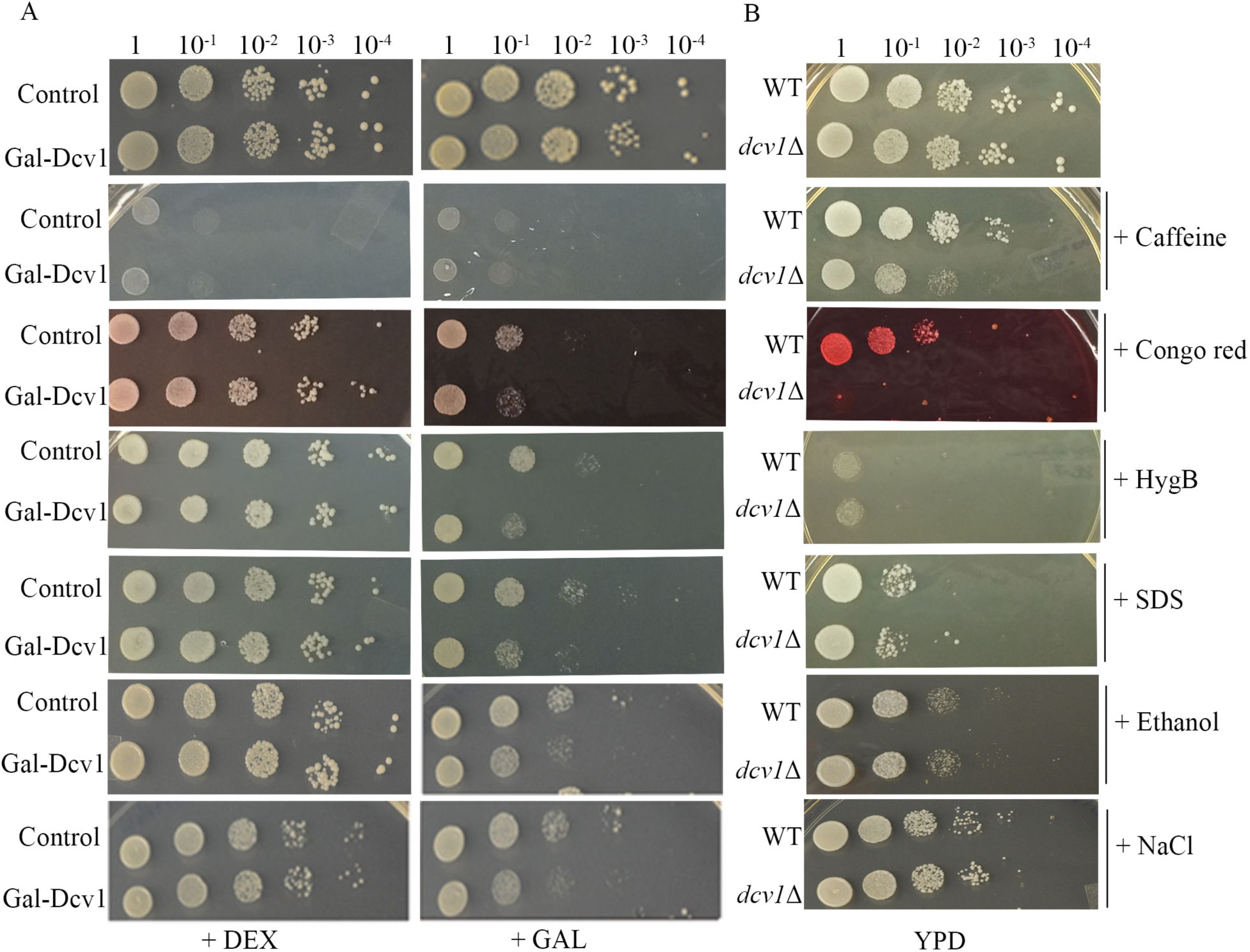
Both overexpression and absence of Dcv1 affect cell integrity. (A) Overexpression of Dcv1 increased sensitivity to all cell wall and PM stressors except caffeine. Log-phase control cells and isogenic cells transformed with Gal1-Dcv1-HA were normalized for cell density and spotted as 10-fold serial dilutions onto plates containing the indicated sugar (2% dextrose or galactose) and the indicated stressors (12mM caffeine, 100 µg/ml Congo Red, 50 µg/ml hygromyocin, 0.001% SDS, 0.4M NaCl, or 4% ethanol). Colonies were allowed to develop for two overnights at 30°C. (B) *dcv1*Δ confers hypersensitivity to Congo Red and increased sensitivity to caffeine. Log-phase WT and *dcv1*Δ cells were tested for sensitivity to the same concentrations of cell wall and PM stressors as described in panel A.

**Figure S3:**
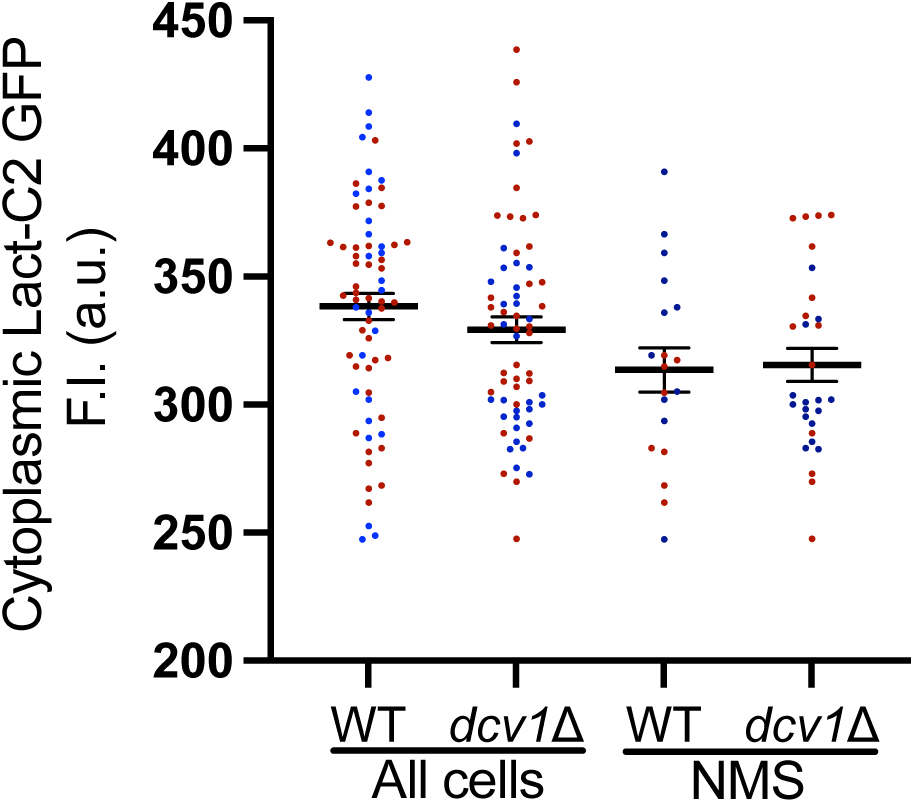
The mean cytoplasmic fluorescence of the PS reporter is comparable in WT and *dcv1*Δ cells. The internal fluorescence of elutriated WT and *dcv1*Δ cells was measured in randomly chosen cells (all cells) and in cells showing no PM signal (NMS) using ImageJ. Data points represent the fluorescent intensities (F.I.) for each category measured in two independent experiments, indicated by color. Lines and error bars represent mean ± s.e.m. (WT: all cells = 338.4 ± 5.1, NMS = 313.5 ± 13.1; *dcv1*Δ: all cells = 329.3 ± 4.9, NMS = 315.5 ± 6.5. n: all cells *≥* 66; NMS *≥* 19. p: all cells = 0.2; NMS = 0.85).

**Figure S4:**
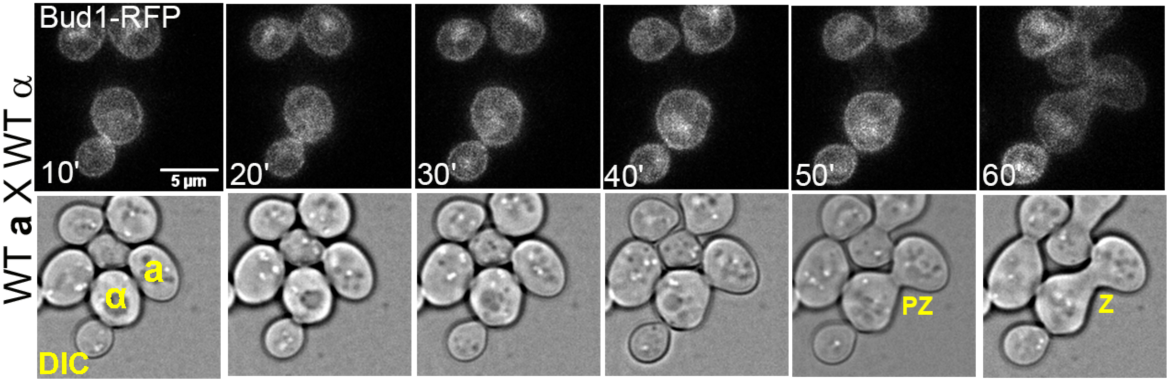
Use of Bud1-RFP to mark *MAT*α cells and to detect zygote formation. Log-phase *MAT***a** cells were mixed with an equal number of *MAT*α cells expressing Bud1-RFP and imaged until fusion of the mating partners. The appearance of RFP in the *MAT***a** partner cells is indicative of cell fusion. The mating partners are labeled **a** and α in the DIC images; PZ, prezygote; Z, zygote.

**Supplementary Table 1:**
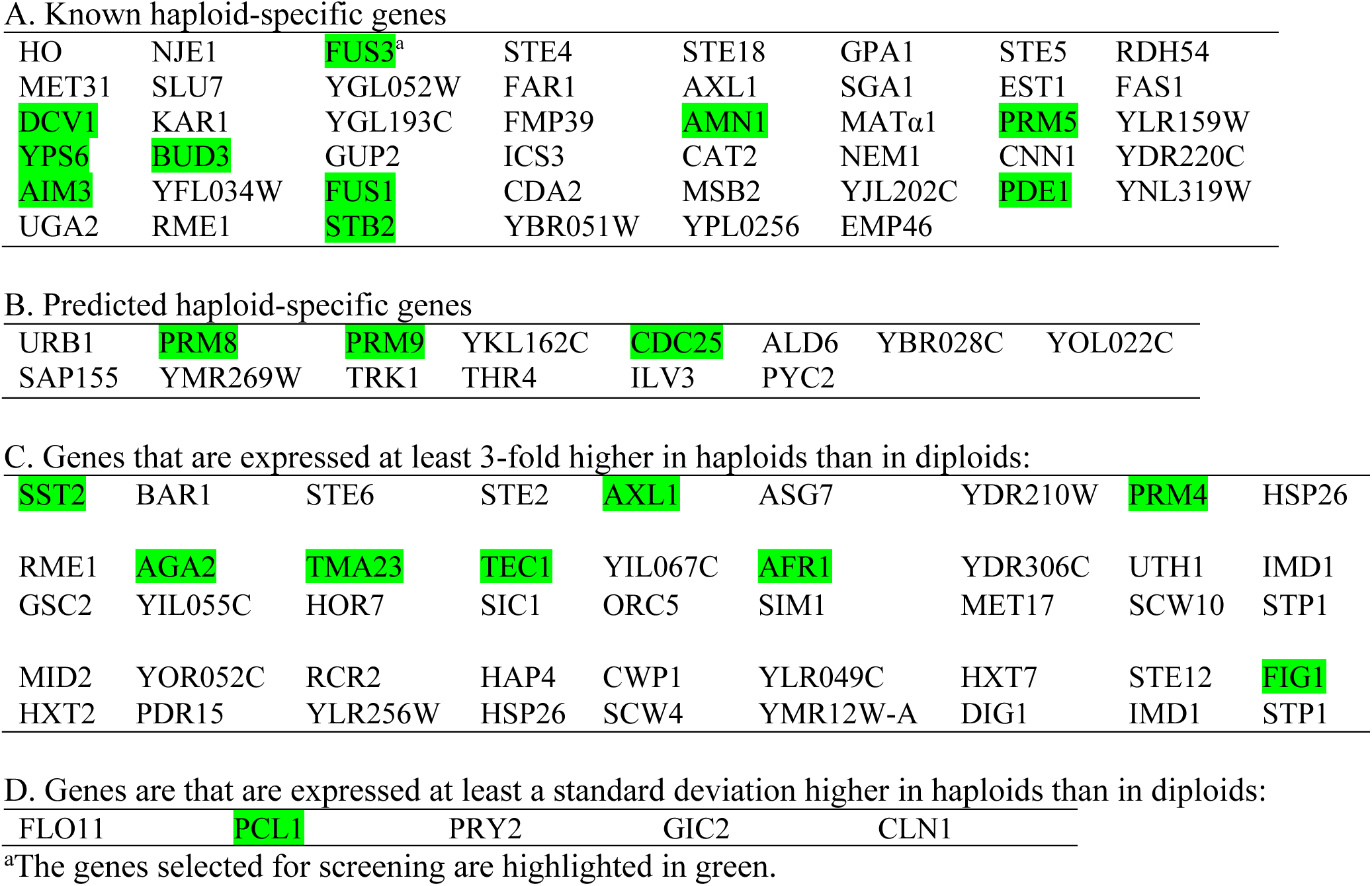
List of haploid-specific genes.

**Supplementary Table 2:**
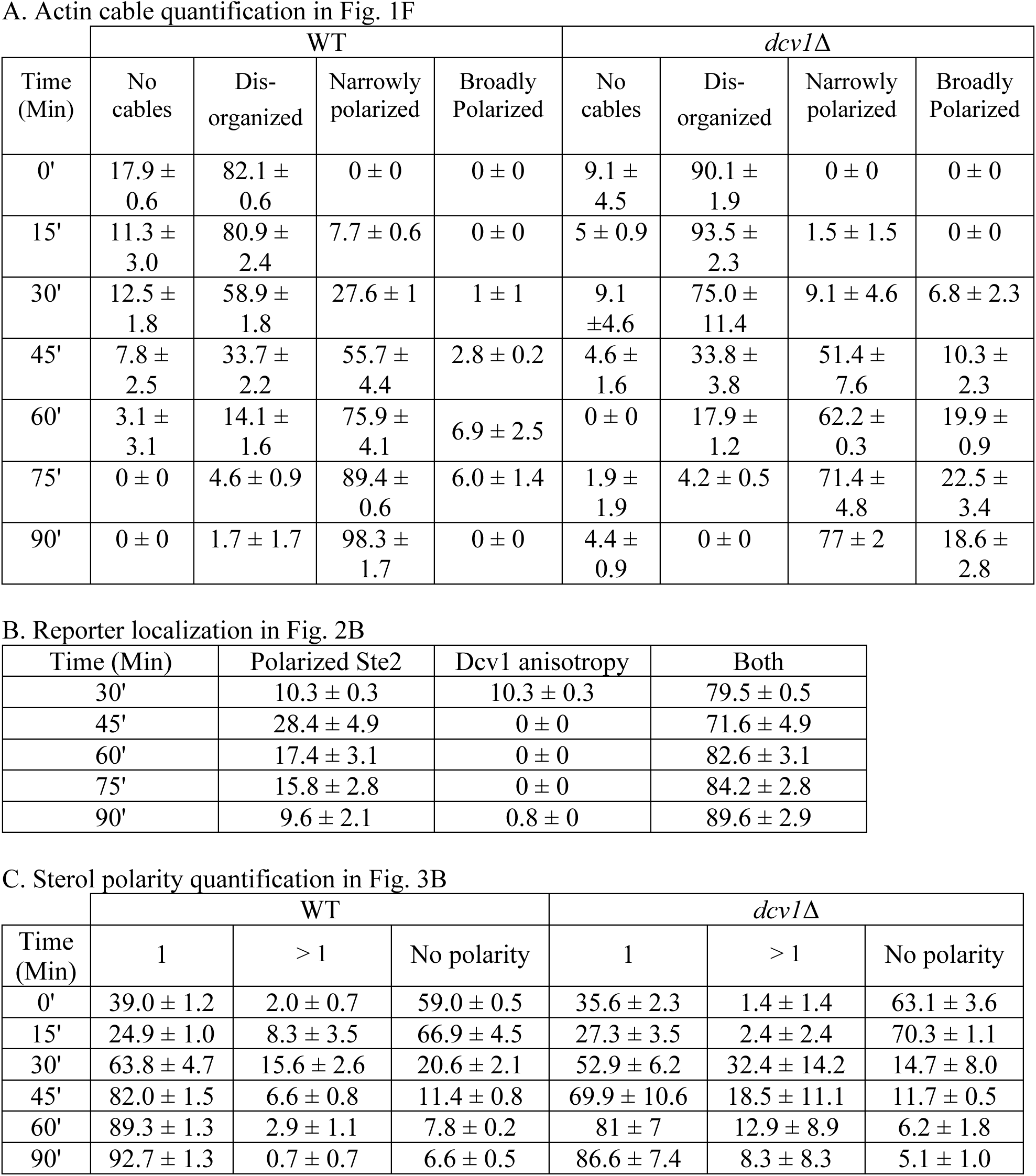
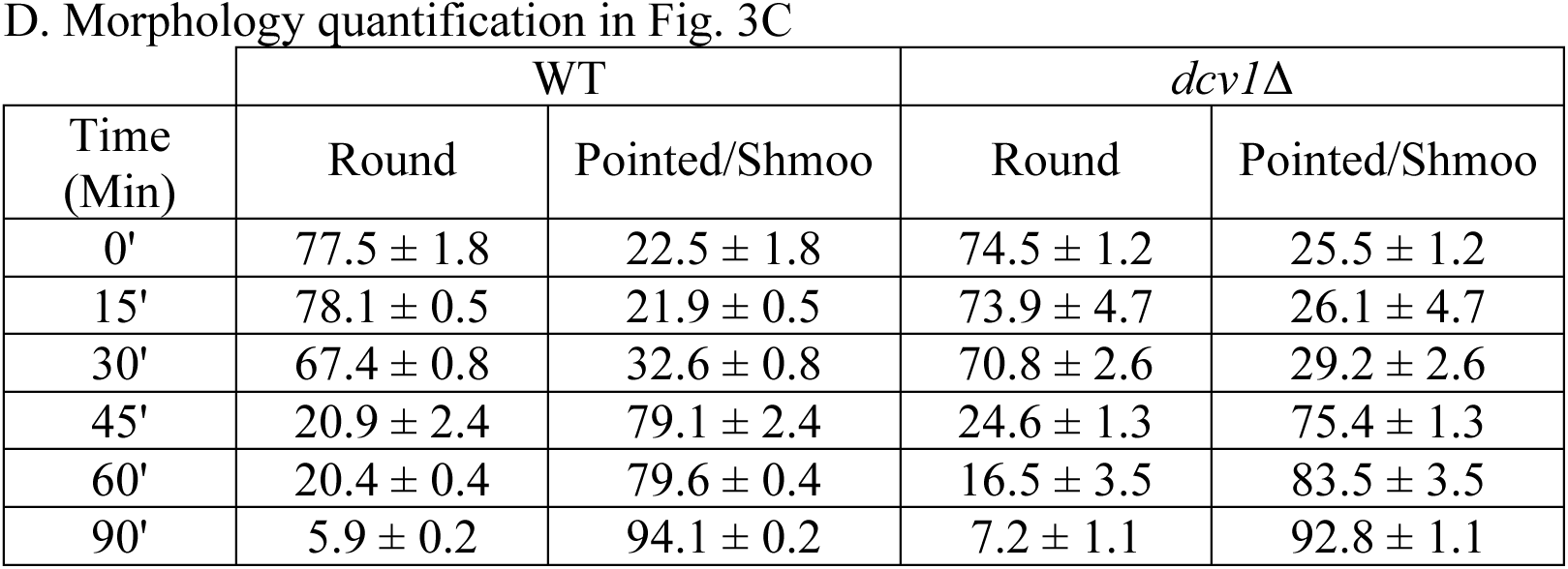
Mean ± s.e.m. values.

**Supplementary Table S3:**
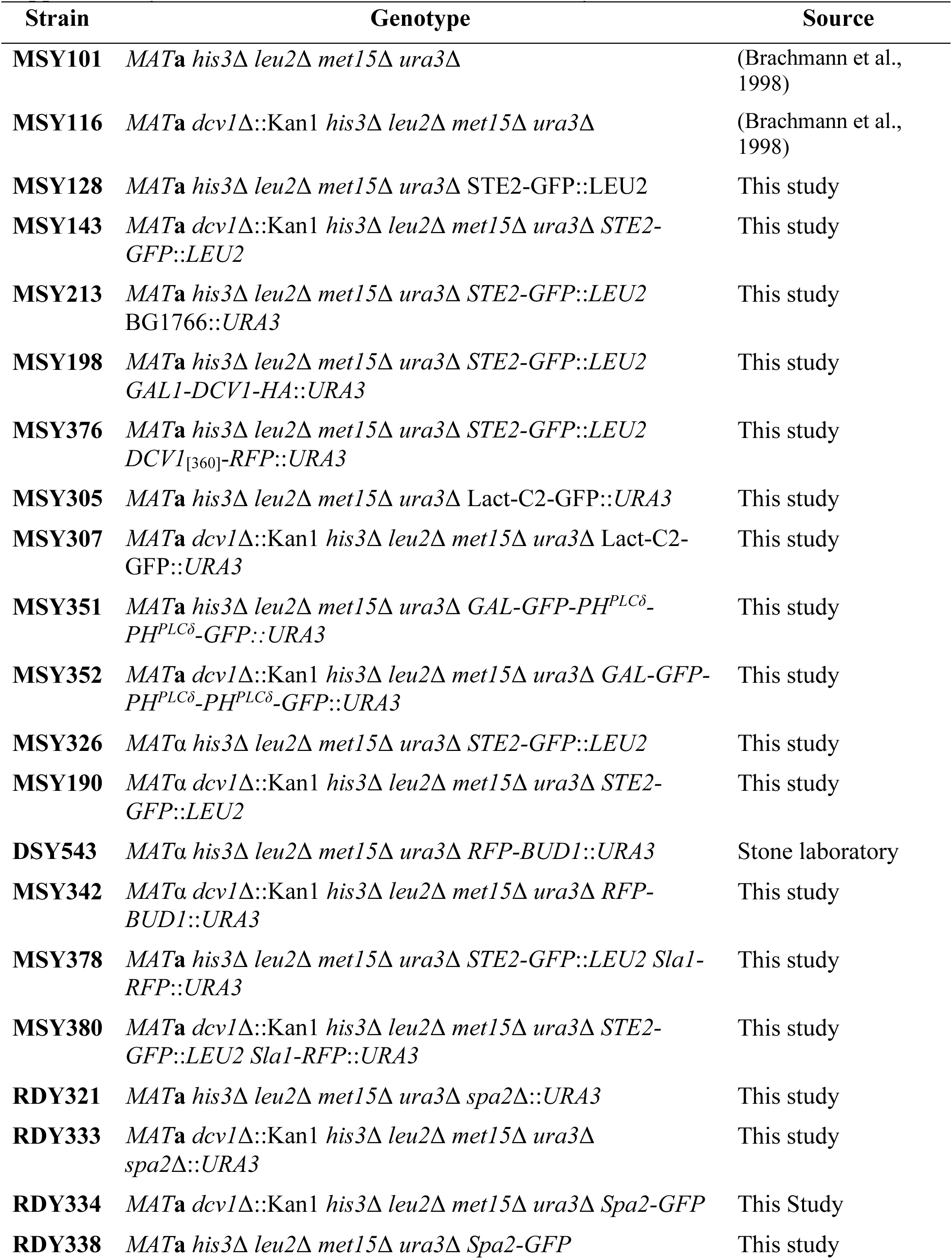
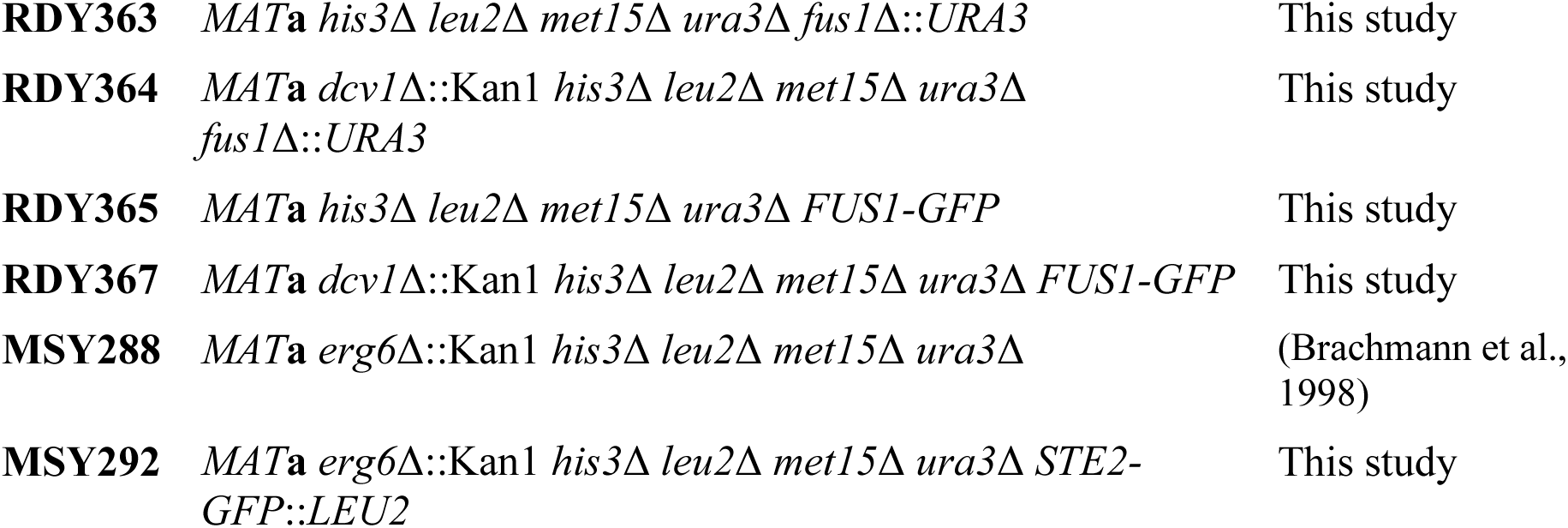
Yeast strains used in this study.

**Supplementary Table S4:** Plasmids used in this study.

| **Plasmid no.** | **Plasmid/Protein expressed** | **Marker/Type** | **Source** |
| --- | --- | --- | --- |
| **YCplac33** | - | URA3/CEN | (Gietz and Akio, 1988) |
| **YIplac211** | - | URA3/CEN | (Gietz and Akio, 1988) |
| **ZWE159** | YCplac22/Gal1-Gal10 | TRP1/CEN | Stone laboratory |
| **LHP1921** | STE2<sup>1-419</sup>-GFP | LEU2/INT | (Dunn et al., 2004) |
| **MSB19** | GAL1-HO | URA3/2μm | (Herskowitz and Jensen, 1991) |
| **MSB20** | BG1805/GAL1-DCV1-19KDA | URA3/2μm | Horizon discovery ORF collection; #YSC3867-202326631 |
| **MSB45** | BG1766 | URA3/2μm | Elizabeth Grayhack |
| **MSB104** | YCplac33/DCV1<sub>[360]</sub>-RFP | URA3/CEN | This study |
| **MSB59** | P406/GAL-GFP-PH<sup>PLCδ</sup>-PH<sup>PLCδ</sup>-GFP | URA3/2μm | Robert Arkowitz |
| **MSB67** | YIplac211/Gal1-Gal10 | URA3/INT | This study |
| **MSB56** | LACT-C2-GFP-p416 | URA3/2μm | (Yeung et al., 2008); Addgene #22853 |
| **MSB68** | YIplac211/GAL1-LACT-C2-GFP | URA3/INT | This study |
| **DSB405** | RFP-BUD1 | URA3/INT | Stone laboratory |
| **XWB087** | YIplac211-Sla1<sup>2491-3732</sup>-RFP | URA3/INT | (Wang et al., 2019) |
| **RDB151** | pRS406/SPA2-GFP | URA3/INT | (Arkowitz and Lowe, 1997) |
| **DSB379** | pRS316-FUS1-GFP | URA3/CEN | Stone laboratory |

## REFERENCES

Aguilar, P. S., Heiman, M. G., Walther, T. C., Engel, A., Schwudke, D., Gushwa, N., Kurzchalia, T. and Walter, P. (2010). Structure of sterol aliphatic chains affects yeast cell shape and cell fusion during mating. Proc. Natl. Acad. Sci. USA 107, 4170–4175.

Alvarez, F. J., Douglas, L. M., Rosebrock, A. and Konopka, J. B. (2008). The Sur7 Protein Regulates Plasma Membrane Organization and Prevents Intracellular Cell Wall Growth in Candida albicans. Mol. Biol. Cell 19, 5214–5225.

Arkowitz, R. A. and Lowe, N. (1997). A Small Conserved Domain in the Yeast Spa2p Is Necessary and Sufficient for Its Polarized Localization. J. Cell Biol. 138, 17–36.

Ausubel, F. M., Brent, R., Kingston, R. E., Moore, D. D., Seidman, J. G., Smith,J. A. and Struhl, K. (1994). Current Protocols in Molecular Biology. John Wiley and Sons, Inc

Ayscough, K. R. and Drubin, D. G. (1998). A role for the yeast actin cytoskeleton in pheromone receptor clustering and signalling. Curr. Biol. 8, 927–930.

Bagnat, M. and Simons, K. (2002). Cell surface polarization during yeast mating. Proc. Natl. Acad. Sci. USA 99, 14183–14188.

Banavar, S. P., Gomez, C., Trogdon, M., Petzold, L. R., Yi, T. M. and Campàs, O. (2018). Mechanical feedback coordinates cell wall expansion and assembly in yeast mating morphogenesis. *PLoS Comp*. Biol. 14, 1–19.

Beh, C. T. and Rine, J. (2004). A role for yeast oxysterol-binding protein homologs in endocytosis and in the maintenance of intracellular sterol-lipid distribution. J. Cell Sci. 117, 2983–2996.

Bidlingmaier, S. and Snyder, M. (2004). Regulation of polarized growth initiation and termination cycles by the polarisome and Cdc42 regulators. J. Cell Biol. 164, 207–218.

Brachmann, C. B., Davies, A., Cost, G. J., Caputo, E., Li, J., Hieter, P. and Boeke, J. D. (1998). Designer deletion strains derived from Saccharomyces cerevisiae S288C: A useful set of strains and plasmids for PCR-mediated gene disruption and other applications. Yeast 14, 115–132.

Butty, A. C., Pryciak, P. M., Huang, L. S., Herskowitz, I. and Peter, M. (1998). The role of Far1p in linking the heterotrimeric G protein to polarity establishment proteins during yeast mating. Science 282, 1511–1516.

Clemente-Ramos, J. Á., Martín-García, R., Sharifmoghadam, M. R., Konorni, M., Osumi, M. and Valdivieso, M. H. (2009). The tetraspan protein Dni1p is required for correct membrane organization and cell wall remodelling during mating in Schizosaccharomyces pombe. Mol. Microbiol. 73, 695–709.

Costanzo, M., VanderSluis, B., Koch, E. N., Baryshnikova, A., Pons, C., Tan, G., Wang, W., Usaj, M., Hanchard, J., Lee, S. D., et al. (2016). A global genetic interaction network maps a wiring diagram of cellular function. Science 353, 1381–1395.

Curto, M. Á., Moro, S., Yanguas, F., Gutiérrez-González, C. and Valdivieso, M. H. (2018). The ancient claudin Dni2 facilitates yeast cell fusion by compartmentalizing Dni1 into a membrane subdomain. Cell. Mol. Life Sci. 75, 1687–1706.

Dobbelaere, J. and Barral, Y. (2004). Spatial coordination of cytokinetic events by compartmentalization of the cell cortex. Science 305, 393–396.

Dunn, R., Klos, D. A., Adler, A. S. and Hicke, L. (2004). The C2 domain of the Rsp5 ubiquitin ligase binds membrane phosphoinositides and directs ubiquitination of endosomal cargo. J. Cell Biol. 165, 135–144.

Erdman, S., Lin, L., Malczynski, M. and Snyder, M. (1998). Pheromone-regulated Genes Required for Yeast Mating Differentiation. J. Cell Biol. 140, 461–483

Fairn, G. D., Hermansson, M., Somerharju, P. and Grinstein, S. (2011). Phosphatidylserine is polarized and required for proper Cdc42 localization and for development of cell polarity. Nat. Cell Biol. 13, 1424–1430.

Garrenton, L. S., Stefan, C. J., McMurray, M. A., Emr, S. D. and Thorner, J. (2010). Pheromone-induced anisotropy in yeast plasma membrane phosphatidylinositol-4,5-bisphosphate distribution is required for MAPK signaling. Proc. Natl. Acad. Sci. USA 107, 11805–11810.

Gehrung, S. and Snyder, M. (1990). The SPA2 gene of Saccharomyces cerevisiae is important for pheromone-induced morphogenesis and efficient mating. J. Cell Biol. 111, 1451–1464.

Georgiev, A. G., Sullivan, D. P., Kersting, M. C., Dittman, J. S., Beh, C. T. and Menon, A. K. (2011). Osh proteins regulate membrane sterol organization but are not required for sterol movement between the ER and PM. Traffic 12, 1341–1355.

Gietz, R. D. and Akio, S. (1988). New yeast-Escherichia coli shuttle vectors constructed with in vitro mutagenized yeast genes lacking six-base pair restriction sites. Gene 74, 527–534.

Goode, B. L., Eskin, J. A. and Wendland, B. (2015). Actin and endocytosis in budding yeast. Genetics 199, 315–358.

Guthrie, C. and Fink, G. R. (2015). Guide to Yeast Genetics and Molecular Biology. San Diego, CA: Academic Press.

He, B., Xi, F., Zhang, X., Zhang, J. and Guo, W. (2007). Exo70 interacts with phospholipids and mediates the targeting of the exocyst to the plasma membrane. EMBO J. 26, 4053– 4065.

Herskowitz, I. and Jensen, R. E. (1991). Putting the HO gene to work: Practical uses for mating-type switching. Methods Enzymol. 194, 132–146.

Howard, J. P., Hutton, J. L., Olson, J. M. and Payne, G. S. (2002). Sla1p serves as the targeting signal recognition factor for NPFX(1,2)D-mediated endocytosis. J. Cell Biol. 157, 315–326.

Ismael, A., Tian, W., Waszczak, N., Wang, X., Cao, Y., Suchkov, D., Bar, E., Metodiev, M. V, Liang, J., Arkowitz, R. A., et al. (2016). Gbeta promotes pheromone receptor polarization and yeast chemotropism by inhibiting receptor phosphorylation. Sci. Signal. 9, 1–17.

Jin, H., McCaffery, J. M. and Grote, E. (2008). Ergosterol promotes pheromone signaling and plasma membrane fusion in mating yeast. J. Cell Biol. 180, 813–826.

Kurihara, L. J., Beh, C. T., Latterich, M., Schekman, R. and Rose, M. D. (1994). Nuclear Congression and Membrane Fusion: Two Distinct Events in the Yeast Karyogamy Pathway. J. Cell Biol. 126, 911–923.

Lingaraju, A., Long, T. M., Wang, Y., Austin, J. R. and Turner, J. R. (2015). Conceptual barriers to understanding physical barriers. Semin. Cell. Dev. Biol. 42, 13–21.

Martin, D. C., Kim, H., Mackin, N. A., Maldonado-Báez, L., Evangelista, C. C., Beaudry, V. G., Dudgeon, D. D., Naiman, D. Q., Erdman, S. E. and Cunningham, K. W. (2011). New regulators of a high affinity Ca2+ influx system revealed through a genome-wide screen in yeast. J. Biol. Chem. 286, 10744–10754.

Merlini, L., Dudin, O. and Martin, S. G. (2013). Mate and fuse: how yeast cells do it. Open Biol. 3, 1–13.

Morioka, S., Shigemori, T., Hara, K., Morisaka, H., Kuroda, K. and Ueda, M. (2013). Effect of sterol composition on the activity of the yeast G-protein-coupled receptor Ste2. Appl. Microbiol. Biotechnol. 97, 4013–4020.

Nelson, B., Parsons, A. B., Evangelista, M., Schaefer, K., Kennedy, K., Ritchie, S., Petryshen, T. L. and Boone, C. (2004). Fus1p Interacts with Components of the Hog1p Mitogen-Activated Protein Kinase and Cdc42p Morphogenesis Signaling Pathways to Control Cell Fusion during Yeast Mating. Genetics 166, 67–77.

Nern, A. and Arkowitz, R. A. (1998). A GTP-exchange factor required for cell orientation. Nature 391, 195–198.

Nern, A. and Arkowitz, R. A. (1999). A Cdc24p-Far1p-Gbeta-gamma Protein Complex Required for Yeast Orientation during Mating. J. Cell Biol. 144, 1187–1202.

Olivera-Couto, A. and Aguilar, P. S. (2012). Eisosomes and plasma membrane organization. Mol. Genet. Genomics 287, 607–620.

Omnus, D. J., Cadou, A., Thomas, F. B., Bader, J. M., Soh, N., Chung, G. H. C., Vaughan, A. N. and Stefan, C. J. (2020). A heat-sensitive Osh protein controls PI4P polarity. BMC Biol. 18, 1–21.

Pringle, J. R., Preston, R. A., Adams, A. E. M., Stearns, T., Drubin, D. G., Haarer, B. K. and Jones, E. W. (1989). Fluorescence Microscopy Methods for Yeast. Methods Cell. Biol. 31, 357–435.

Proszynski, T. J., Klemm, R., Bagnat, M., Gaus, K. and Simons, K. (2006). Plasma membrane polarization during mating in yeast cells. J. Cell Biol. 173, 861–866.

Pruyne, D. and Bretscher, A. (2000). Polarization of cell growth in yeast. J. Cell Sci. 113, 571– 585.

Sartorel, E., Unlu, C., Jose, M., Aurélie, M. L., Meca, J., Sibarita, J. B. and McCusker, D. (2018). Phosphatidylserine and GTPase activation control Cdc42 nanoclustering to counter dissipative diffusion. Mol. Biol. Cell 29, 1299–1310.

Sherman, F., Fink, G. R. and Hicks, J. B. (ed.) (1986). Laboratory Course Manual For Methods in Yeast Genetics. *Cold Spring Harbor, NY*: *Cold Spring Harbor Laboratory Press*.

Shi, C., Kaminskyj, S., Caldwell, S. and Loewen, M. C. (2007). A role for a complex between activated G protein-coupled receptors in yeast cellular mating. Proc. Natl. Acad. Sci. USA 104, 5395–5400.

Stefan, C.J., Audhya, A., Emr, S.D. (2002). The yeast synaptojanin-like proteins control the cellular distribution of phosphatidylinositol (4,5)-bisphosphate. Mol. Biol. Cell. 13, 542–557.

Suchkov, D. V., DeFlorio, R., Draper, E., Ismael, A., Sukumar, M., Arkowitz, R. and Stone, D. E. (2010). Polarization of the Yeast Pheromone Receptor Requires Its Internalization but Not Actin-dependent Secretion. Mol. Biol. Cell 21, 1737–1752.

Tiedje, C., Holland, D. G., Just, U. and Höfken, T. (2007). Proteins involved in sterol synthesis interact with Ste20 and regulate cell polarity. J. Cell Sci. 120, 3613–3624.

Trueheart, J., Boeke, J. D. and Fink, G. R. (1987). Two genes required for cell fusion during yeast conjugation: evidence for a pheromone-induced surface protein. Mol. Cell Biol. 7, 2316–2328.

Valdez-Taubas, J. and Pelham, H. R. B. (2003). Slow Diffusion of Proteins in the Yeast Plasma Membrane Allows Polarity to Be Maintained by Endocytic Cycling. Curr. Biol. 13, 1636–1640.

Vasanji, A., Ghosh, P. K., Graham, L. M., Eppell, S. J. and Fox, P. L. (2004). Polarization of plasma membrane microviscosity during endothelial cell migration. Dev. Cell 6, 29–41.

Walther, T. C., Brickner, J. H., Aguilar, P. S., Bernales, S., Pantoja, C. and Walter, P. (2006). Eisosomes mark static sites of endocytosis. Nature 439, 998–1003.

Wang, X., Tian, W., Banh, B. T., Statler, B. M., Liang, J. and Stone, D. E. (2019). Mating yeast cells use an intrinsic polarity site to assemble a pheromone-gradient tracking machine. J. Cell Biol. 218, 3730–3752.

Yeung, T., Gilbert, G. E., Shi, J., Silvius, J., Kapus, A. and Grinstein, S. (2008). Membrane Phosphatidylserine Regulates Surface Charge and Protein Localization. Science 319, 210– 213.

